# Pre-chlorotic Mg^2+^ deficiency impairs guard-cell PSII and leaf gas exchange in Vicia faba

**DOI:** 10.64898/2026.10.07.757235

**Authors:** Andrea Wübben, Evelien Franck, Bastian Leander Franzisky, Muhammad Ishfaq, Alexander Erban, Joachim Kilian, Edda von Roepenack-Lahaye, Joachim Kopka, Mark Stahl, Sebastien Carpentier, Christoph-Martin Geilfus, Katja Kampmann

## Abstract

Magnesium deficiency impairs photosynthesis and transpiration before visible chlorosis, but whether guard cells themselves are affected at this early stage remains unresolved. We combined whole-leaf gas exchange and chlorophyll fluorescence with cell-specific fluorescence measurements of individual guard cell pairs, together with elemental analysis, metabolite profiling, adenosine triphosphate quantification, and proteome profiling of guard cell-enriched material in hydroponically grown *Vicia faba*. Magnesium deficiency reduced magnesium concentrations in both whole leaves and guard cell-enriched fractions. Under red light, magnesium-deficient leaves showed reduced effective quantum yield of photosystem II, CO_2_ assimilation, transpiration, and stomatal conductance, whereas intercellular CO_2_ concentration remained unchanged. Individual guard cell pairs likewise showed reduced effective quantum yield of photosystem II, demonstrating that the early deficiency response extended to guard cells. Electron transport rate and non-photochemical quenching in guard cells were not significantly altered. Only a small set of metabolites in guard cell-enriched material, mainly fatty acids and citrate, responded to magnesium deficiency, whereas ATP abundance and light-induced proton extrusion remained unchanged. In contrast, the guard cell-enriched proteome showed higher abundance of photosynthesis- and translation-related proteins despite reduced guard-cell PSII efficiency. Together, these findings identify guard cells as an early responsive cell type under magnesium deficiency and show that guard-cell photochemistry is impaired before visible chlorosis, alongside reduced whole-leaf carbon gain and water loss.

## 1 Introduction

Beside impairments in various physiological processes, magnesium (Mg^2+^) deficiency negatively affects photosynthesis, transpiration (E) and stomatal conductance (gs) (Houmani et al., 2024; Li et al., 2024; Liu et al., 2025; Parisa et al., 2026; Xie et al., 2024). So far, these effects are commonly explored through bulk-leaf analysis and shown as a consequence from mesophyll-driven processes like impaired CO_2_ fixation, accumulation of photosynthetic end-products and the development of reactive oxygen species (ROS) which may damage chloroplasts (H. Marschner and Cakmak, 1989) and therefore alter the leaf pigment status. In summary, this results in the decrease of photosynthesis (Cakmak and Kirkby, 2008). As Mg^2+^ deficiency reduces mesophyll CO fixation it potentially alters internal or apoplastic CO_2_ and sugar signals, which may influence stomatal aperture. Regarding this, Zörb et al. (2022) reported that Mg^2+^ limited E might be related to apoplastic CO_2_ concentrations. And further, that CO_2_ consumption of red light-illuminated mesophyll cells reduces leaf apoplastic CO_2_ concentrations, which might lead into stomatal widening to facilitate CO uptake (Roelfsema et al., 2002). Zörb et al. (2022) highlights that Mg^2+^ nutrition and E could be linked through these mesophyll-driven CO_2_ concentrations or sugar signals. More recently, apoplastic sugars were identified as mesophyll-derived signals that promote red-light-induced stomatal opening by regulating guard-cell H^+^-ATPase activity and ion transport in Arabidopsis and *Vicia faba* (Zait et al., 2025). Overall, stomatal function is widely considered to be predominantly regulated by mesophyll-derived signals, based on products of photosynthesis or electron transport, the tissue redox state, metabolites transported in the transpiration stream, and vapor-phase ionic and/or electrical signals (Lawson et al., 2014).

Guard cells (GCs) harbor chloroplasts, and there is a consensus, that electron transport and enzymes of the Calvin-Benson cycle are functional. However, it remains controversial if GC photosynthesis contributes in stomatal regulation (Lemonnier and Lawson, 2024). Tominaga et al. (2001) suggests that GC chloroplasts provide ATP required for plasma membrane H⁺-ATPase activity and stomatal opening. This is consistent with the evidence that red light induces photosynthesis-dependent phosphorylation of the H^+^-ATPases in GCs (Ando and Kinoshita, 2018), suggesting that osmoregulation in GCs supports stomatal opening. Azoulay-Shemer et al. (2015) reported that reduced chloroplasts in GCs leads into reduced gs and assimilation rate, although stomata still respond to CO_2_ alterations. Recent reviews emphasize that GCs are metabolically active and harbor multiple metabolic pathways, including photosynthetic electron transport, the Calvin–Benson cycle, starch, sucrose and malate metabolism, as well as mitochondrial respiration, all of which have been implicated in stomatal function (Lawson and Matthews, 2020; Lemonnier and Lawson, 2024). More recently, guard-cell-specific manipulation showed that changes in GC metabolism can alter stomatal behavior (Sun et al., 2026). However, it is unknown whether Mg^2+^ deficiency reaches GC before visible chlorosis and affects their photochemical function. Given that Mg²⁺ affects the above listed photochemical and metabolic processes at the whole-leaf level, such a depletion would be expected to restrict the same processes in GCs. We therefore hypothesize that early Mg^2+^ deficiency also affects GC photochemistry and metabolism. We therefore compared Mg^2+^-sufficient and Mg^2+^-deficient *Vicia faba* plants at the pre-chlorotic stage and contrasted whole-leaf responses with GC-associated responses, using the oldest leaves after the cotyledons (L2) and a red light-treatment. Mg^2+^ concentrations were determined in bulk leaf tissue as well as in GC enriched tissue to determine whether Mg^2+^ depletion reaches this cell type.

Fluorescence measurements were executed to investigate effective quantum yield of PSII (Y(II)), quantum yield of non-photochemical quenching (Y(NPQ)), inner carbon content (Ci), as well as CO_2_ assimilation (*A*), *E* and g_s_ in mesophilic tissue. To mirror this analysis for GC responses we measured Y(II), Y(NPQ) and the electron transport rate (ETR) on isolated GCs. The pH was additionally compared between epidermal peel discs and leaf discs. Furthermore, the ATP, metabolome and proteome status of GC-enriched material was investigated.

## Material and Methods

### Experimental design

Plants were grown hydroponically under two Mg^2+^ treatments: an adequately supplied control (named “+Mg” via 0.3 mM MgSO_4_) and a Mg^2+^-deficient treatment (named “-Mg”, no MgSO_4_). All analyses were performed on the oldest fully expanded leaves after the cotyledons (leaf (L) 2, abbreviated L2) between 14 and 18 days after transfer to hydroponics, when -Mg plants had reached a Mg²⁺-deficient status but visible chlorosis was still absent or not yet pronounced (Fig. 1). To separate bulk-leaf from GC-related responses, L2 leaves, epidermal peels and GC-enriched fractions obtained via the blender method were analyzed in parallel. The leaves represented mainly mesophyll-dominated bulk-leaf responses, epidermal peels largely excluded mesophyll effects, and blender-derived GC-enriched fractions provided sufficient material for biochemical and molecular analyses. The intact L2 leaves were used for spectral reflectance measurements (n = 6), whole-leaf Mg^2+^ analysis (n = 9) and gas exchange and chlorophyll fluorescence measurements (n = 4). Epidermal peels were used for single-GC chlorophyll fluorescence measurements, with one GC pair from each independent plant representing one biological replicate (n = 7 for +Mg, n = 8 for -Mg), and for extracellular pH-related measurements using epidermal peel discs (n = 6). For comparison with the peel-based extracellular pH response, leaf discs were analyzed in parallel to determine the pH change induced by bulk-leaf tissue in the external medium (n = 5). GC-enriched material obtained with the blender method was used for GC Mg^2+^ analysis, ATP measurements, metabolite- and proteome-profiling. For GC Mg^2+^ analysis, one hydroponic pot containing five plants, corresponding to ten L2-leaves, represented one biological replicate (n = 6 for +Mg, n = 4 for -Mg). For ATP (epidermal peels n = 6, leaf discs n = 5), metabolome (pre-dawn n = 38, post-dawn n = 4-7) and proteome analyses (n = 5), L2 leaves from three hydroponic pots, corresponding to 15 plants and 30 L2 leaves, were pooled into one biological replicate. Here, proteomic and metabolomic profiling was performed on GC-enriched material rather than on whole-leaf tissue. This design allowed Mg^2+^ deficiency responses at the whole-leaf level to be compared with responses occurring specifically in GC-enriched material.

**Figure 1:**
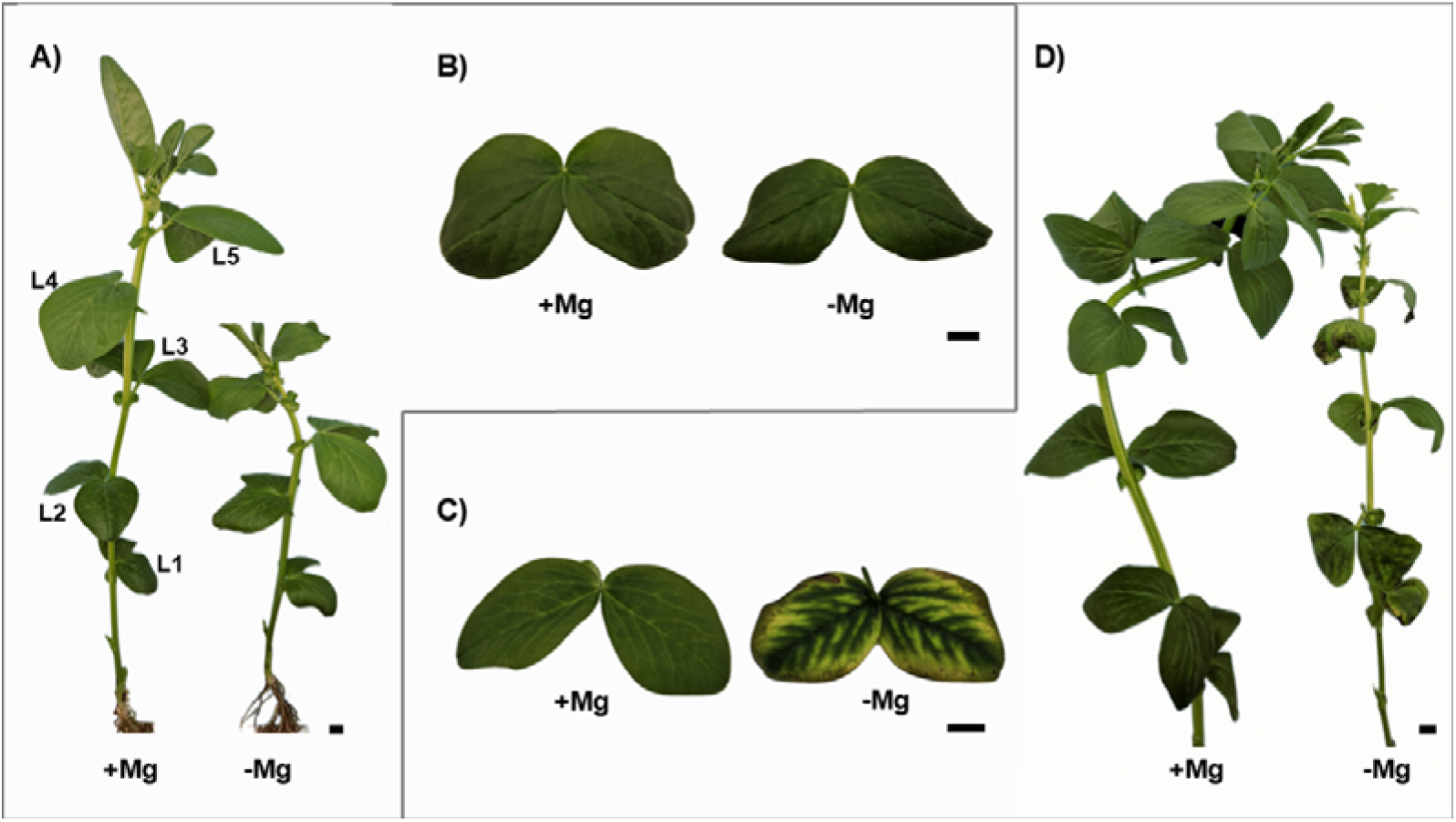
*V. faba* plants (A) and D)) and fully expanded leaves (L2, B) and C)), 16 (A), B)) and 23 (C), D)) days after transferred to hydroponic conditions. +Mg received a well supply of Mg (0.3 mM). No Mg was supplied in the - Mg group. In **A)** the positions of full expanded leaf pairs are schematically shown on +Mg, starting with the cotyledons at leaf position one (L1). Scale bars at lower right indicate 1 cm.

### Plant cultivation

Seeds of *V. faba* (cv. Allison) were provided by Norddeutsche Pflanzenzucht (NPZ, Holtsee, Germany) and were germinated on quartz sand (0.4 – 0.8 mm). After 9 days, plants were transferred to a hydroponic system, with five plants placed into one 5 l pot, containing nutrient solution (NL). To acclimate the plants, the initial NL contained one-quarter of the full-strength formulation. After four days, plants were transferred to the full-strength NL (0.1 mM KH_2_PO_4_, 1.5 mM K_2_SO_4_, 0.5 mM KCl, 2.0 mM Ca(NO_3_)_2_, 60 μM Fe-EDTA, 50 μM Na_2_O_3_Si, 10 μM H_3_BO_4_, 2.0 μM MnSO_4_, 0.5 μM ZnSO_4_, 0.2 μM CuSO_4_, 0.05 μM NiCl_2_ and 0.05 μM (NH_4_)_6_Mo_7_O_24_), which was renewed every seven days. +Mg received adequate supply of magnesium (Mg^2+^) as MgSO_4_ (0.3 mM) whereas -Mg received no MgSO_4_ during hydroponic cultivation. Sulfur was supplied in sufficient amounts in both treatments to meet the nutritional requirements of field bean.

### Guard cell enrichment

#### Epidermal peels

Following Cornish and Zeevaart (1986) and Geilfus et al. (2018), epidermal peels were taken from the abaxial surface of the L2 leaves. To remove adhering mesophyll cells, peels were placed into 5 ml reaction tubes filled with deionized water and treated for 45 s with an ultrasonic homogenizer (50 W, 20% cycle and 50% amplitude; UP200St ultrasonic processor with sonotrode S26d2, Heilscher Ultrasonics GmbH, Teltow, Germany). To ensure stomata were closed, peels were floated on distilled water following Schwarz et al (1990) until measurements began.

#### Blender method

GC were enriched following Bauer et al. (2013) with minor modifications. Excised leaves were separated from the middle veins and blended for 60 s at highest level (Braun, JB 3060 700 W, Neu-Isenburg, Germany) with 200 ml deionized water containing crushed ice. GCs were harvested on a 210 µm pore wide nylon mesh and blended again for 60 s at highest level with 200 ml deionized water containing crushed ice. Intact GC with adhering fragments of lysed pavement cells were then harvested with the specified mesh and placed in 2 ml reaction tubes. Both methods were earlier verified by means of viability stains (Geilfus et al., 2018; Geilfus et al., 2015).

### Spectral reflectance measurements

For determination of modified chlorophyll absorption in reflectance 1 (MCARI1), the carotenoid-sensitive indices 1 (CRI1) and the normalized difference vegetation index (NDVI), the PolyPen RP 410 (PSI, Czech Republic) was used (Lee at al. 2024). Leaves were clamped in the PolyPen leaf clip. Reflectance spectra were recorded at four measurement points per leaf side (adaxial and abaxial) and averaged. Material was taken at day 16 after plants were transferred to hydroponic cultivation.

### Magnesium measurement

Mg^2+^ concentrations were measured using inductively coupled plasma optical emission spectrometry (ICP-OES, SPECTRO ARCOS, SPECTRO Analytical Instruments GmbH, Kleve, Germany). Sample preparation varied depending on tissue type. To determine Mg^2+^ concentrations in the leaves, L2 leaves were dried at 60°C, weighed and digested according to VDLUFA (2008) with minor alterations: grinding was omitted. Approximately 0.25 g dry matter was digested with 3 ml H_2_O, 7 ml HNO_3_ and 2 ml H_2_O_2_. To determine Mg^2+^ in GC-enriched samples, GC were collected as described before (‘blender method’) and freeze-dried for 48 h after storage at −80°C. Mg was analyzed following VDLUFA (VDLUFA, 2008) with minor alterations. Grinding was omitted, the dry matter weight was approximately 0.02 g and 0.9 ml H_2_O, 2.1 ml HNO_3_ and 0.6 ml H_2_O_2_ were added. 6 ml H_2_O were added to the digestion vessels and sample tubes were placed inside. After digestion, samples were rinsed up to 15 ml with H_2_O and then filtered.

### Gas exchange measurements on leaves

For gas exchange measurements on intact L2 leaves, *E*, *A*, *g_s_*, intercellular CO_2_ concentration (*C_i_*), Y(II) and Y(NPQ) were measured, using a CIRAS-4 Portable Photosynthesis System (PP Systems, Amesbury, MA, USA). Plants were taken 2 h post-dawn and were dark adapted for 30 min. Afterwards, measurements were carried out on intact L2 leaves under 1000 µmol m^−2^ s^−1^ red light above canopy at 22°C, 450 µmol mol^−1^ CO and 70% relative humidity corresponding to ambient conditions. Data points were collected at steady-state conditions after 10 min. Saturation pulses (6000 µmol m^−2^ s^−1^ intensity and 1 s duration) were applied every 15 s over 10 min.

### Microscopy-PAM measurements of chlorophyll fluorescence in guard cells

Chlorophyll fluorescence parameters of individual GC pairs were measured using a Microscopy Pulse-Amplitude-Modulated fluorometer (Microscopy-PAM, Heinz Walz GmbH, Effeltrich, Germany). Epidermal peels from L2 leaves were prepared as described above and were kept in distilled water to prevent curling. To reduce tissue irritation, peels were taken 1.5 h after peeling and mounted on microscope slides in 50 mM KCl solution. Measurements were then performed on single GC pairs under low-intensity red light (6 µmol m^−2^ s^−1^), providing a defined dawn-like light stimulus while avoiding strong activation of surrounding tissue. The microscope focus and aperture were adjusted to the selected GC pair to obtain cell-type-specific fluorescence data. Saturation pulses were applied at the beginning of the experiment, after 60 s and after 10 min to determine Y(II), ETR and Y(NPQ). To minimize unintended light effects, the bright-field source and auxiliary lamp were covered with green foil.

### Quantification of ATP in GC-enriched leaf material

Plant material from L2 leaves was collected, and GC-enriched material was isolated using the blender method described above. Half of the material was frozen in liquid nitrogen and stored at – 80°C to serve as pre-dawn samples. The other half was incubated in 50 mM KCl solution in petri dishes and illuminated with 50 µmol m^−2^ s^−1^ red light for 2 h210 µm pore widethe material was freeze-dried for 48 h later and weighed (about 10 mg per sample) into 2 ml reaction tubes. GC-enriched samples were then ground twice for one minute at a frequency of 30 s^−1^ using a mixer mill (MM 400, Retsch GmbH, Haan, Germany) containing one 5 mm stainless steel ball. The powder was suspended in 1.5 ml of ultrapure water, kept on ice and vortexed for 30 s to dissolve ATP. Samples were centrifuged afterwards at 14400 g for 10 min at 4°C and frozen in liquid nitrogen for storage. Measurements were performed using the Enliten® ATP assay system. Luminescense was measured at 560 nm for 10 sec per sample using the TECAN Spark® platereader (TECAN Spark, Tecan Trading AG, Männedorf, Switzerland) giving relative light units (RLU). A 7-point external calibration from 10 amol to 10 pmol ATP was performed for absolute quantitation.

### Metabolic and proteomic profiling of GC-enriched material

Using the blender method as described above, GC were enriched from collected L2 material. Half of each sample was immediately frozen in liquid nitrogen, while the other half was incubated in 50 mM KCl solution in petri dishes. Samples were illuminated with 50 µmol m^−2^ s^−1^ red light for 2 h210 µm pore wideground twice for one minute at a frequency of 30 s^−1^ using a mixer mill (MM 400, Retsch GmbH, Haan, Germany) containing one 5 mm stainless steel ball each. For metabolome analysis, soluble metabolites were extracted according to Richter et al. (2019) with modifications. 570 µl MeOH/CH_3_Cl/H_2_O (2.5/1/1, v/v/v) and 30 µl ^13^C_6_-sorbitol (0.2 mg ml^−1^) and the frozen sample were homogenized and thawed for 5 min in a vortex mixer. Samples were then incubated at 70°C for 15 min and 950 rpm and centrifuged at 14.000 rcf for 10 min. 150 µl of the supernatant was transferred into a new reaction tube and sonicated for 5 min in 400 µl MeOH/CH_3_Cl/H_2_O (2.5/1/1, vol/vol/vol), heated at 70°C for 10 min (950 rpm), and centrifuged at 14.000 rcf for 10 min. The remaining sample was re-extracted and sonicated for 5 min with 400 µl MeOH/CH_3_Cl/H_2_O (2.5/1/1, vol/vol/vol), heated at 70°C for 10 min (950 rpm), and centrifuged at 14.000 rcf for 10 min. 500 µl of the supernatant was combined with the first extract of 150µL and subsequently dried in a vacuum concentrator overnight at room temperature. Gas chromatography-electron impact/time-of-flight mass-spectrometry (GasC-EI/TOF-MS)-based metabolite profiling was carried out after Dethloff et al. (2014) in split less mode. Chromatogram-data files were visually monitored, baseline-corrected, and exported in the NetCDF file format, using the ChromaTOF software (version 4.22, LECO, St. Joseph, USA). With the TagFinder software (Luedemann et al., 2008), the GasC-EI/TOF-MS data were converted into a chromatographically aligned numerical table format of mass features, defined by nominal mass (m/z), retention time index and abundance, i.e., peak height of arbitrary units. Abundance data were expressed in arbitrary units, standardized by an internal standard, U^13^C - sorbitol, normalized to dry weight and background-subtracted. Compounds were identified according to Dethloff and co-authors (2014) by matching mass spectral and retention time index reference data of the Golm Metabolome Database, GMD (http://gmd.mpimp-golm-mpg.de/search.aspx) (Hummel et al, 2010, Kopka et al. 2005).

### Proteomic profiling

Proteins of 5 samples per magnesium treatment were extracted. 500 µL of extraction buffer (Table S1) was added to the lyophilized guard cells and vortexed until homogenized. 500 µL of Tris-buffered phenol (pH 8.0) was added and shaken for 10 min. at 4°C, followed by 10 min. of centrifugation at 12000 rpm (g force 8900) at 4°C. The phenolic phase was collected and the extraction process was repeated once more. Afterwards, 5 volumes of ice cold 100 mM ammonium acetate in methanol were added to the phenolic phase, inverted and stored at −20°C overnight. Then, the samples were centrifuged for 1 hour at 13000 rpm (g force 10400) at 4°C. After removing the supernatant, the pellet was rinsed with 2 mL rinsing solution (cold acetone/0.2 % DTT) for 1 hour at −20°C, followed by centrifugation for 30 min. at 13000 rpm at 4°C. After removing the supernatant, the pellet was air dried and resuspended in 50 µL lysis buffer (Table S2) and vortexed shortly. Proteins were quantified using the 2-D Quant kit (Cytiva) according to the manufacturer’s manual and aliquoted per 20 µg proteins. Samples were prepared for liquid chromatography-mass spectrometry (LC-MS) by mixing 5 µg of protein with 10% sodium dodecyl sulfate and 50 mM triethylammonium bicarbonate followed by S-trap protocol. Peptides were analyzed using an Evosep One LC system coupled to a SCIEX ZenoTOF mass spectrometer. Chromatographic separation was performed using the 60 samples per day (60 SPD) Evosep method with a 21-minute gradient. Mass spectrometry data were acquired in SWATH data-independent acquisition (SWATH/DIA) mode. Data were analyzed using Spectronaut v19.5.241126.62635 (Biognosys AG) in directDIA mode (directDIA+ Deep workflow) (Biognosys AG, 2023). Peptide identification was performed in a library-free workflow using dynamic MS1 and MS2 extraction with intensity-based fragment ion selection. Retention time alignment was achieved using source-specific iRT calibration with local non-linear regression, excluding deamidated peptides. Searches assumed trypsin/P specificity with up to two missed cleavages, carbamidomethylation of cysteine as a fixed modification, and methionine oxidation and protein N-terminal acetylation as variable modifications. False discovery rates were controlled at 1% at the PSM, peptide, and protein group levels. Spectronaut-specific q-value filtering was applied at threshold of 0.01 for precursor at the experiment level and 0.05 for proteins at the run level.Quantification was performed at the MS2 level using area-based integration, followed by cross-run normalization using Spectronaut’s automatic strategy without imputation. Protein abundances were derived using mean peptide intensities with Top-N aggregation (1–3 precursors per protein group) and interference correction restricted to identified peptides. Protein annotations and GO terms for *Vicia faba* (Taxon ID 3906) were retrieved from UniProt (https://www.uniprot.org/taxonomy/3906, retrieved on 06/01/2026).

### Quantification of light-induced proton extrusion by guard cells

Epidermal peels were prepared using the epidermal peel method as described above, with minor alterations. Peels were taken 1.5 h pre-dawn and 3.5 mm diameter discs were cut out of the leaf material and peel fragment using a biopsy punch. Discs were incubated in distilled water for 1.5 h. 0.5 h before dawn discs were placed in a PCR-reaction tube each, containing 50 µl of 50 mM KCl and were then illuminated with 50 µmol m^−2^ s^−1^ red for 2 h. After illumination, the incubation solution, containing protons extruded by GC, was collected and stored at −80°C until external pH quantification. Measurements were done using ratiometric fluorometry with the pH-sensitive fluorescent dye Oregon Green 488, following the principle described by Geilfus and Mühling (2011). Samples were excited sequentially at 440 and 495 nm, and fluorescence emission was recorded at approximately 525–535 nm. The fluorescence ratio F_495_/F_440_ was calculated and converted into pH values using a calibration curve. Light-induced proton extrusion was assessed from the acidification of the external incubation medium.

### Statistical analysis

Statistical analyses were performed using GraphPad Prism 10 (GraphPad Software, San Diego, CA, USA). A Shapiro-Wilk test was applied to test for normal distribution. Depending on data distribution, either an unpaired two-tailed t-test with or without Welch’s correction or a two-tailed Mann-Whitney U test was performed. For Mg^2+^ concentrations in leaves (Fig. 2A) an unpaired t-test was applied and for the measurements in GC an unpaired t-test with Welch’s correction was used. In Fig. 3 an unpaired t-test was examined for the comparisons of MCARI1, CRI1 and NDVI for the adaxial side (Fig. 3A-C) and unpaired t-tests with Welch’s correction were performed for the comparisons on the abaxial side (Fig. 3D-F). For gas exchange measurements on leaves (Fig. 4) data was analyzed using an unpaired t-test. Statistical analysis for Microscopy-PAM measurements was performed using an unpaired t-test with Welch’s correction (Fig.5A and 5C) and a two-tailed Mann-Whitney U test (Fig. 5B). For pH analysis, an unpaired t-test (Fig. 6A) or a Mann-Whitney test (Fig. 6B) was applied on the proton concentration, calculated from measured pH values. For ATP measurements Mann-Whitney U tests were performed for pre-dawn and post-treatment measurements (Fig. 6C-D). The GC metabolome was statistically analyzed in R version 4.4.2. Metabolite abundances were log2-transformed. Differences in individual metabolite abundance were assessed using a two-way aligned rank transform (ART) ANOVA (‘ARTool’ package) with the fixed factors Mg²⁺ treatment and the sampling time (pre- or post-dawn) and their interaction. Only metabolites with more than 4 observations in each group were analyzed, resulting in 61 estimable metabolites (Tab. S3). Missing values were handled by available-case analysis without imputation. P-values were adjusted for multiple testing using the Benjamini-Hochberg false discovery rate (FDR) correction. To assess whether the overall metabolite profile separated the two Mg²⁺ treatments, a partial least squares discriminant analysis (PLS-DA, ‘mixOmics’ package) with two components was performed. Because sampling time dominated the overall variance, the PLS-DA was restricted to pre-dawn samples (n = 38). An exploratory PLS-DA including both sampling times showed a strong separation between pre- and post-dawn samples, while the Mg^2+^ treatments largely overlapped. Therefore, treatment effects were further analyzed using pre-dawn samples only. In this analysis, the two Mg^2+^ treatments separated along the first component, with a cross-validated BER of 0.15 (permutation P = 0.005). For this analysis, metabolites with less than 10 % missing values were retained (n = 44) and the few remaining missing values were imputed as the observed minimum minus one on the log2 scale. The abundances were scaled to unit variance. Classification performance was evaluated by repeated five-fold cross-validation (50 repeats) and expressed as the balanced error rate (BER). A permutation test (200 permutations of the treatment labels) was performed to assess the significance of the separation.

**Figure 2:**
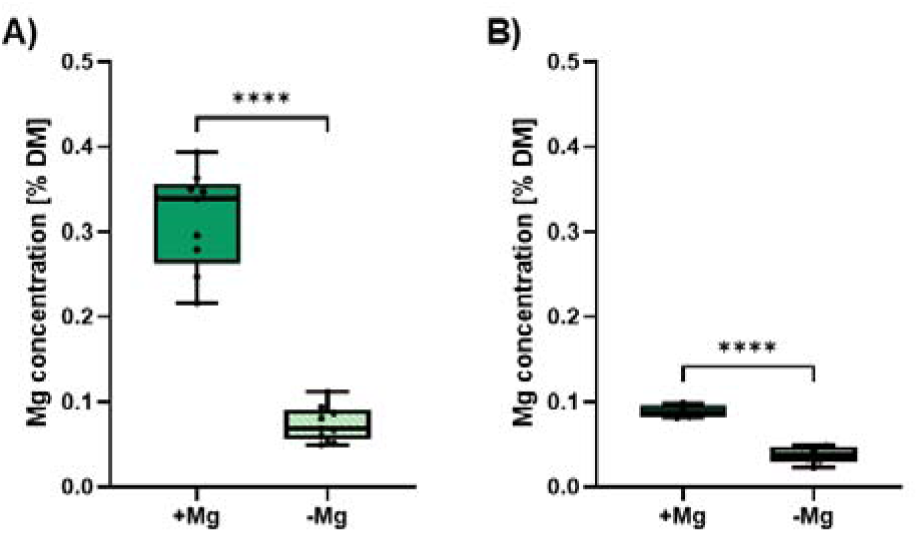
Mg concentration of fully expanded, detached leaves (A)) of L2, and enriched-GC material (B)), 16 days after transferred to hydroponics. +Mg (filled boxes) received a well supply of Mg^2+^ (0.3 mM). No Mg was supplied in -Mg (hatched boxes). Boxplots were generated with GraphPad Prism 10 (GraphPad Software, San Diego, CA, USA) showing the 25 to 75 percentile range. Each data point represents one leaf pair of a single plant (**A)**) or GC-enriched material isolated from 10 L2 leaves (**B)**). For **A)** an unpaired t-test was applied (n=9) and for **B)** an unpaired t-test with Welch’s correlation was used (n=6 for +Mg and n=4 for –Mg). Lines inside the boxes indicate the median, whiskers extend to the minimum and maximum values. Statistical significance is indicated as follows: *P < 0.05, *P < 0.01, ***P < 0.001, ****P < 0.0001.

**Figure 3:**
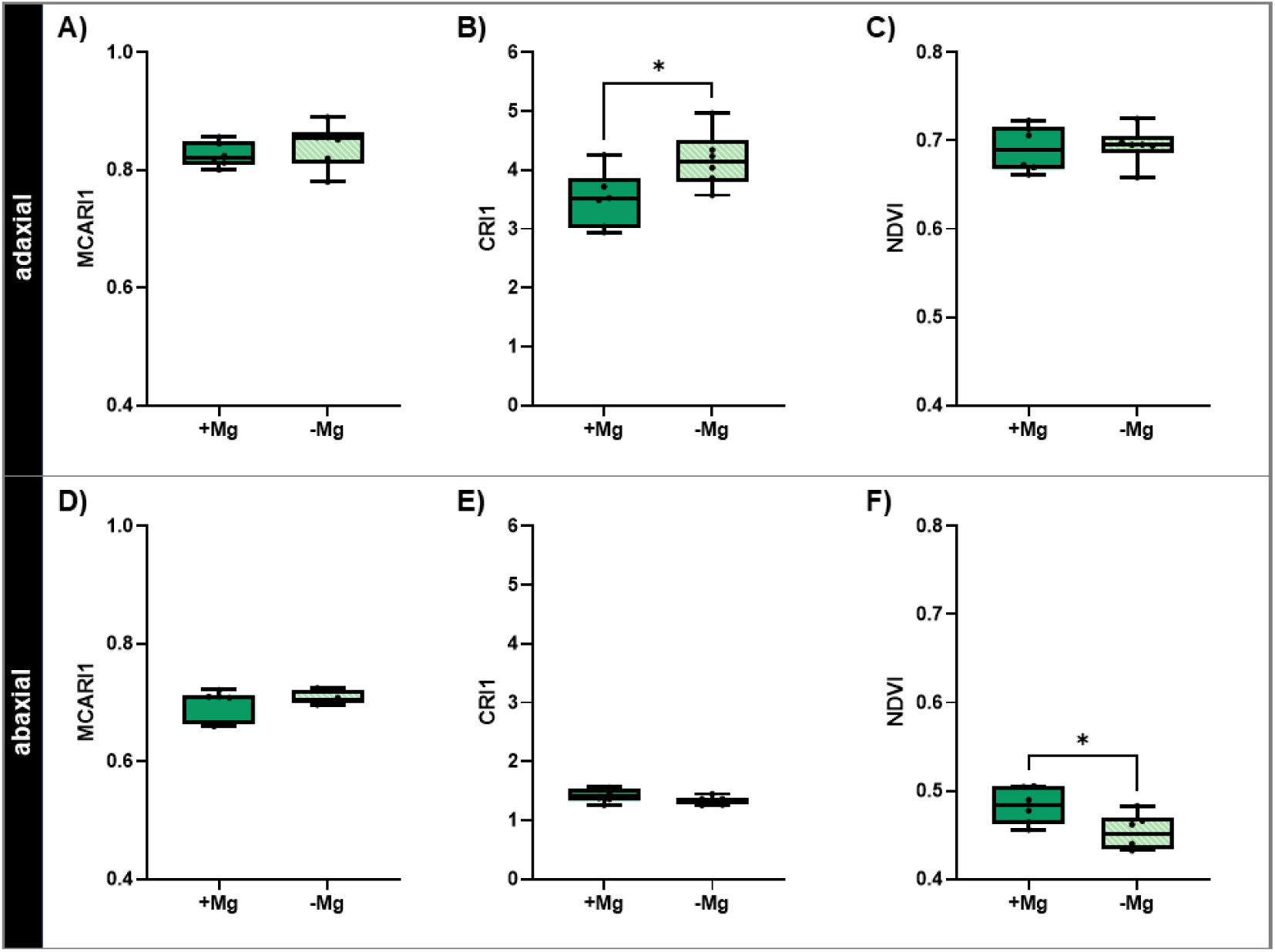
Adaxial and abaxial data of the modified chlorophyll absorption in reflectance 1 (MCARI1) (A) and D)), the carotenoid-sensitive indices 1 (CRI1) (B) and E)), and the normalized difference vegetation index (NDVI) (C) and F)) measured with the PolyPen RP 410 (PSI, Czech Republic) on fully expanded leaves (L 2) 16 days after transferred to hydroponics. +Mg (filled boxes) received a well supply of Mg^2+^ (0 .3 mM). No Mg was supplied in - Mg (hatched boxes). Boxplots were generated with GraphPad Prism 10 (Graph Pad Software, San Diego, CA, USA) showing the 25^th^ to 75^th^ percentile range. Each point represents the mean of four measurement points per leaf side (n=6). An unpaired t-test was examined for the comparisons of MCARI1, CRI1 and NDVI for the adaxial side (**A)-C)**) and unpaired t-tests with Welch’s correlation were performed for the comparisons on the abaxial side (**D)-F)**). Lines inside the boxes indicate the median, whiskers extend to the minimum and maximum values. Statistical significance is indicated as follows: *P < 0.05, **P < 0.01, *** P < 0.001.

**Figure 4:**
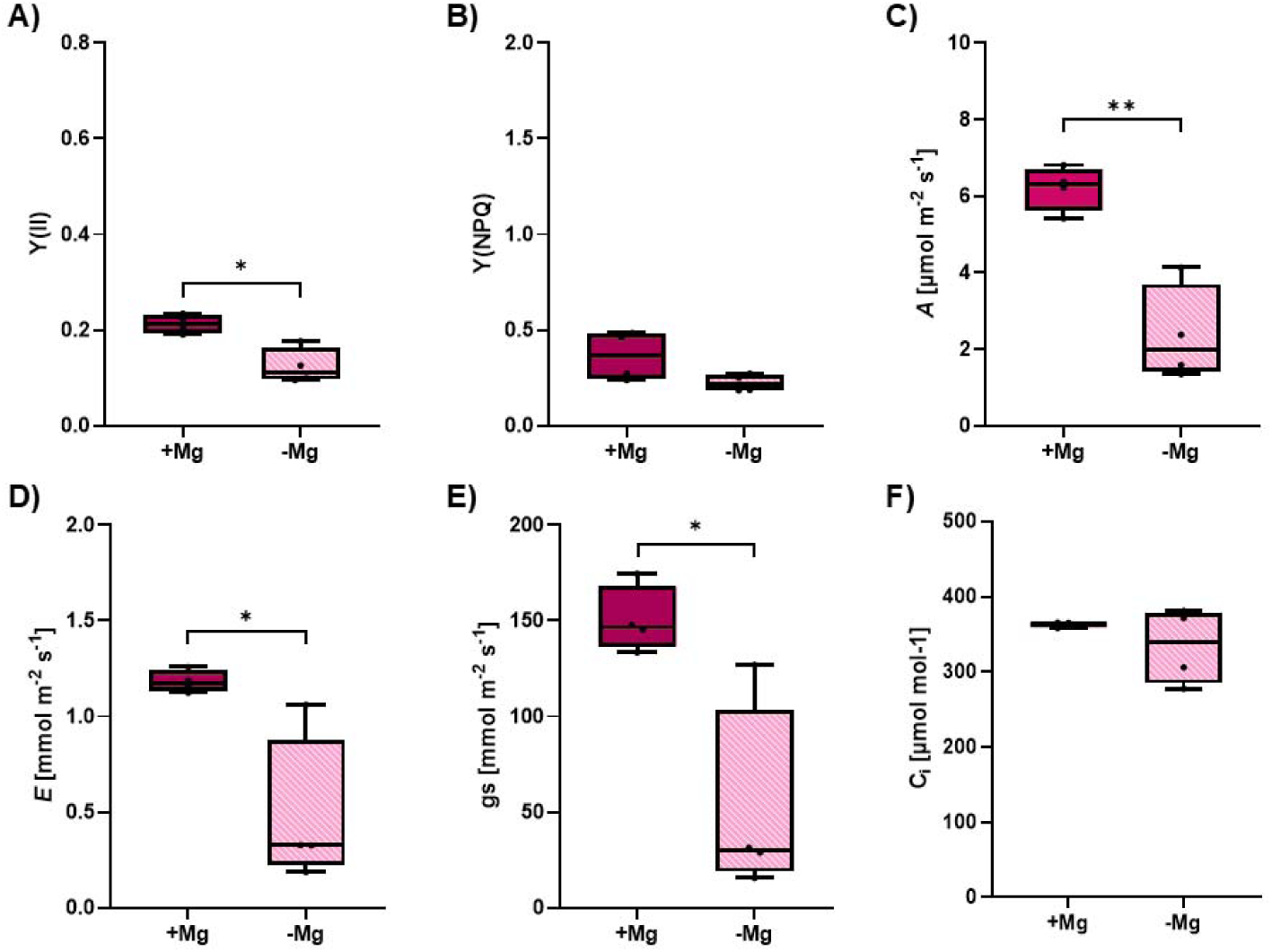
Effective quantum yield of photosystem II (Y(II), A)); quantum yield of non-photochemical quenching (Y(NPQ), B)); net CO_2_ assimilation (A, C)); Transpiration (E, D)); stomatal conductance (g_s_, E)) and inner carbon content (C_i_, F)) of full expanded leaves (L2) of intact plants, two weeks after transferred to hydroponics. +Mg (filled boxes) received a well supply of Mg (0 .3 mM) in a hydroponic system, whereas no Mg was supplied in - Mg (patterned boxes). Plants were taken 2 h after sunset, were dark adapted for 30 min and treated with 1000 µmol m^−2^s^−1^ red light for 10 min. Data shows the end points after 10 min of light treatments. Measurements were performed using a CIRAS-4 Portable Photosynthesis System (PP Systems, Amesbury, MA, USA). Boxplots were generated with GraphPad Prism 10 (GraphPad Software, San Diego, CA, USA) showing the 25^th^ to 75^th^ percentile range. All parameters were analyzed, using an unpaired t-test Each data point represents the measurement of a single leaf (n=4). Lines inside the boxes indicate the median, whiskers extend to the minimum and maximum values. Statistical significance is indicated as follows: *P < 0 .05, **P < 0.01, *** P < 0.001, ****P < 0.0001.

**Figure 5:**
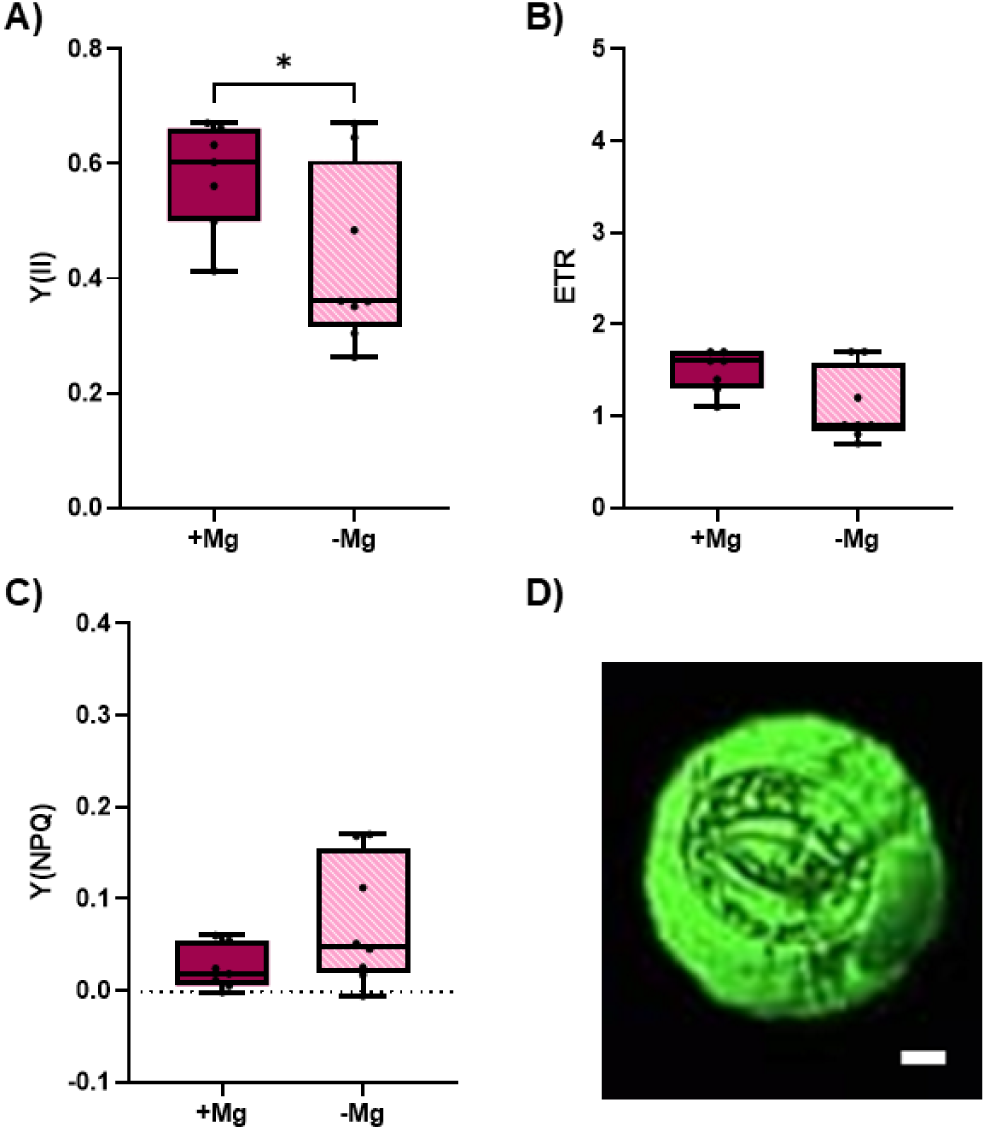
Effective quantum yield of photosystem II (Y(II)) (A)), electron transport rate (ETR) (B)) and quantum yield of non-photochemical quenching (Y(NPQ)) of non-regulated energy dissipation in photosystem II (Y(NO)) (C)) of a single pair of abaxial GCs. +Mg (filled boxes) received a well supply of Mg (0 .3 mM MgSO_4_) in a hydroponic system, whereas no Mg was supplied in - Mg (patterned boxes). Measurements were carried out using a Microscopy Pulse-Amplitude-Modulated fluorometer (Microscopy-PAM, Heinz Walz GmbH, Effeltrich, Germany) by focusing on a single pair of GCs of an epidermal peel. Peels from full expanded leaves (L2) were taken and treated with 6 µmol m^−2^s^−1^ red light for 10 min. Boxplots were gen erated with GraphPad Prism 10 (GraphPad Software, San Diego, CA, USA) showing the 25^th^ to 75^th^ percentile range. Each data point represents the measurement of one GC pair (n=7 for +Mg, n=8 for –Mg). An unpaired t-test with Welch’s correlation was applied for **A)** and **B)** and a Mann-Whitney U test for **C)**. Lines inside the boxes indicate the median, whiskers extend to the minimum and maximum values. Statistical significance i s indicated as follows: *P < 0.05, **P < 0.01, ***P < 0.001. (**D)**) shows a single pair of GCs. Surrounding signal was reduced by adjusting the microscope aperture to the guard cells. Scale indicates 10 µm.

**Figure 6:**
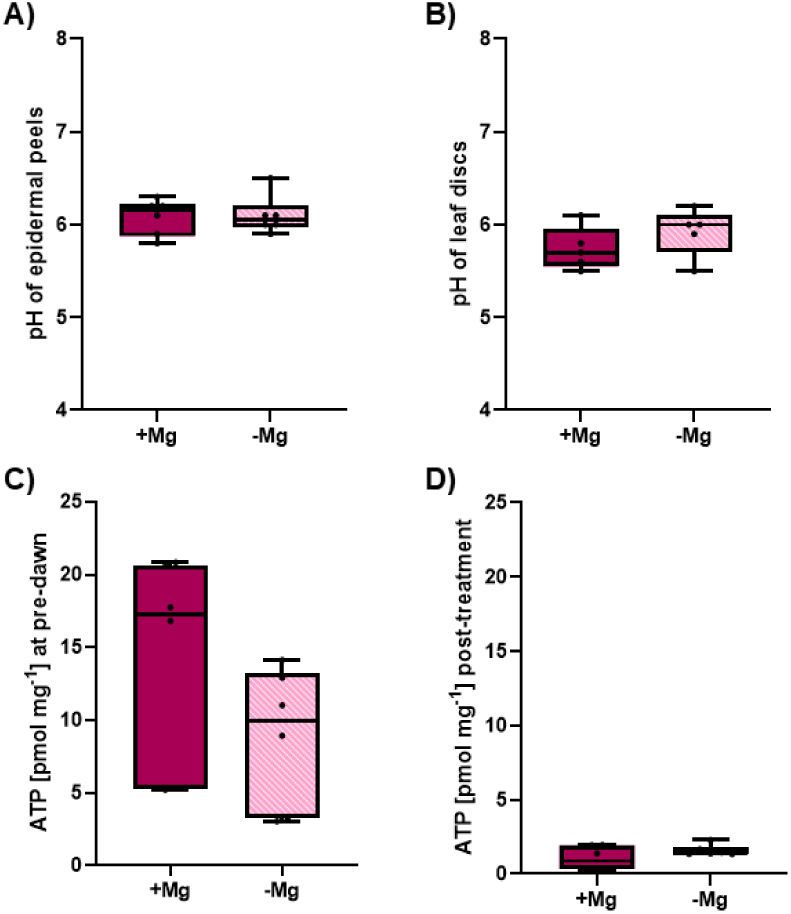
External, i.e. apoplastic, pH measurements and adenosine triphosphate (ATP) concentration in GC-enriched material. +Mg (filled boxes) received a well supply of Mg (0 .3 mM MgSO_4_) in a hydroponic system, whereas no Mg was supplied in -Mg (patterned boxes). For pH measurements outside of GC, discs from an epidermal peel (**A)**) or a leaf (**B)**) were treated with 6 µmol m^−2^s^−1^ red light for 2 h after 1 .5 h incubation time. For ATP measurements GCs were isolated 2 h before dawn using the blender method, where pre-dawn samples were taken (**C)**). 0 .5 h before dawn other half of the samples were treated with 50 µmol m^−2^s^−1^ red light for 2 h to serve as post-treatment samples (**D)**). For p H measurements an unpaired t-test was performed for **A)** and a Mann-Whitney U test for **B)** based on the proton concentration, calculated from measured p H values. For ATP Mann-Whitney U tests applied for **C)** and **D).** Graphics were generated with Graph Pad Prism 10 (Graph Pad Software, San Diego, CA, USA) showing the 25^th^ to 75^th^ percentile range. Each data point represents the measuring point of one disc (n=6 for **A)** and n=5 for **B)**) or a pooled leaf sample, taken from three hydroponic pots (up to 15 leaf samples for **C)** and **D)**, n=6). Lines inside the boxes indicate the median, whiskers extend to the minimum and maximum values. Statistical significance is indicated as follows: *P < 0.05, **P < 0.01, ***P < 0.001.

Analysis of the proteomic profile was performed in R version 4.2.3. Partial Least Square Discriminant Analysis (PLS-DA) (‘mixOmics’ package) with one component was performed to separate samples by Mg^2+^ treatment. Proteins were ranked by Variable Importance in Projection (VIP) scores and a breakpoint was identified using elbow method (‘segmented’ package) (Fig. S1). Proteins with VIP score higher than the breakpoint were analyzed using Gene Ontology (GO) enrichment (‘topGO’ package) using a node size of 5, the weight01 algorithm, Fisher statistic (α = 0.01) with Benjamini-Hochberg FDR correction and 626 nodes tested. Background used were all identified proteins from the +Mg and – Mg samples.

## 3 Results

### 3.1 Pre-chlorotic Mg deficiency reduces guard cell Mg concentration and significantly affects abaxial NDVI and adaxial CRI1

Measurements were conducted on the first fully expanded leaves (L2) above the cotyledons of *V. faba* plants 16 days after they were transferred to a hydroponic system (Fig. 1A). Mg^2+^ deficient L2 leaves did not show visible deficiency symptoms compared to the L2 leaves of the Mg-sufficient treatment (Fig. 1B). By day 23, Mg^2+^-deficient plants exhibited severe interveinal chlorosis with necrotic patches on fully expanded leaves, while leaves of sufficiently Mg^2+^-supplied plants retained their dark green appearance (Fig. 1C, D). Therefore, the L2 leaf pair at 16 days after transfer was selected for subsequent analyses, as it represents an early stage prior to the onset of severe visual symptoms.

The Mg^2+^ concentration of detached fully expanded leaves displayed a significantly lower concentration in the Mg^2+^-deficient treatment (Fig. 2A, P < 0.0001), with a concentration of 0.08% dry matter (DM) (± 0.007) and an approximately 4-fold higher concentration of 0.31 % DM (± 0.019) in the Mg^2+^-sufficient treatment. Additionally, the magnesium concentration of the GC material was significantly lower in the Mg^2+^-deficient treatment, as shown by an unpaired two-tailed t-test (Fig. 2B, P < 0.0001). This treatment had a mean value of 0.04% DM (± 0.004), whereas the Mg^2+^-sufficient treatment had a Mg^2+^ concentration of 0.09% DM (± 0.004).

The Modified Chlorophyll Absorption in Reflectance Index (MCARI1), the Carotenoid Reflectance Index (CRI1), as well as the Normalized Difference Vegetation Index (NDVI) are shown for the adaxial side of the fully expanded leaves of *V. faba* (Fig. 3 A-C) and for the abaxial side of the same leaves (Fig. 3 D-F). MCARI1 was not significantly different between the Mg^2+^-deficient and the Mg^2+^-sufficient treatment, whether adaxial, or abaxial. However, the Mg^2+^-deficient treatment had a significantly higher adaxial CRI1 (P = 0.035) with a value of 4.17 (± 0.19, dimensionless reflectance index). For the abaxial side, no significant differences between the treatments were found. . The NDVI was significantly lower in the Mg^2+^-deficient treatment compared to the Mg^2+^ sufficient treatment for the abaxial side (P = 0.029) with a value of 0.45 (± 0.008) for the Mg^2+^-deficient treatment and 0.48 (± 0.008) for the Mg-sufficient treatment. Also, there was no significant difference regarding the NDVI for the adaxial side.

### 3.2 Mg deficiency reduces photosynthesis under red light in fully expanded leaves

The Y(II) of the fully expanded leaves was significantly lower in the Mg^2+^-deficient treatment (Fig. 4A, P = 0.006). The Mg^2+^-deficient treatment showed a Y(II) of 0.12 (± 0.01), whereas the Mg-sufficient treatment reached values of 0.21 (± 0.01). Moreover, the Mg^2+^-deficient treatment had a generally lower Y(NPQ) (Fig. 4B) with 0.22 (± 0.02) compared to the Mg^2+^-sufficient treatment with 0.36 (± 0.05). Concerning the gas exchange measurements, a significantly lower *A* was found for the Mg^2+^-deficient treatment (Fig. 4C, P = 0.0015) with 2.36 µmol m^−2^ s^−1^ (± 0.49), compared to the Mg^2+^-sufficient treatment with an *A* of 6.20 µmol m^−2^ s^−1^ (± 0.22). Furthermore, the Mg^2+^-deficient treatment showed a significantly lower *E* (Fig. 4D, P = 0.012) with 0.48 mmol m^−2^ s^−1^ (± 0.15), while the Mg^2+^-sufficient treatment reached a *E* of 1.18 mmol m^−2^ s^−1^ (± 0.02). With 50.66 mmol m^−2^ s^−1^ (± 19.86), the g_s_ is significantly reduced in the Mg^2+^-deficient treatment (Fig. 4E, P = 0.029) compared to the Mg^2+^-sufficient control with 150.29 mmol m^−2^ s^−1^ (± 6.73). The Ci did not significantly differ between the Mg^2+^ treatments (Fig. 4F, P = 0.3), with the Mg^2+^-deficient treatment reaching a value of 334.02 µmol mol^−1^ (± 25.19), while the Mg^2+^-sufficient treatment showed a Ci of 362.52 µmol mol^−1^ (± 1.8).

### 3.3 Photosystem II efficiency in guard cells is impaired under red light and Mg deficiency

Single GCs were analysed with PAM-fluorometry to assess photosynthetic variables (Fig. 5D). A significantly lower Y(II) of the Mg^2+^-deficient treatment with 0.43 (± 0.05) compared to the Mg^2+^-sufficient treatment with 0.58 (± 0.04) was revealed by an unpaired t-test with Welch’s correction (Fig. 5A, P = 0.043). There were no significant effects of the Mg^2+^-treatment on the electron transport rate (Fig. 5B) or the Y(NPQ) of the GCS (Fig. 5C). However, the Y(NPQ) was generally higher under Mg^2+^-deficient conditions with 0.08 (± 0.03) compared to the Mg^2+^-sufficient treatment with a value of 0.02 (± 0.01).

### 3.4 Mg deficiency alters guard cell protein accumulation

The pH was examined in discs of epidermal peels (Fig. 6A) as well as in leaf discs (Fig. 6B) of both Mg^2+^ treatments. There were no significant differences between the Mg^2+^-sufficient and the Mg^2+^-deficient treatment, neither in the peel nor in the leaf. External pH did not differ between Mg2+ treatments in epidermal peels or leaf discs. There was only a tendency towards a higher pH of the Mg^2+^-deficient treatment with 5.92 (± 0.12) in the leaf compared to the Mg^2+^-sufficient treatment with 5.74 (± 0.1).

Furthermore, ATP concentration was analyzed in GC-enriched material at pre-dawn (Fig. 6C) and post-dawn (Fig. 6D). ATP concentration did not differ significantly between Mg^2+^ treatments at either sampling time. However, ATP was numerically (but not siginficantly) lower under Mg^2+^ deficiency at pre-dawn and numerically higher at post-dawn. An ART-based two-way ANOVA with FDR-correction was performed to assess differences in GC metabolite concentration at pre- and post-dawn. 41 of the 61 estimable metabolites differed significantly between the two time points (Fig. 7) with the majority showing lower concentrations post-dawn. A Mg^2+^ effect was observed for 8 metabolites, including fatty acids and citrate. These concentrations were reduced under Mg^2+^ deficiency.

**Figure 7:**
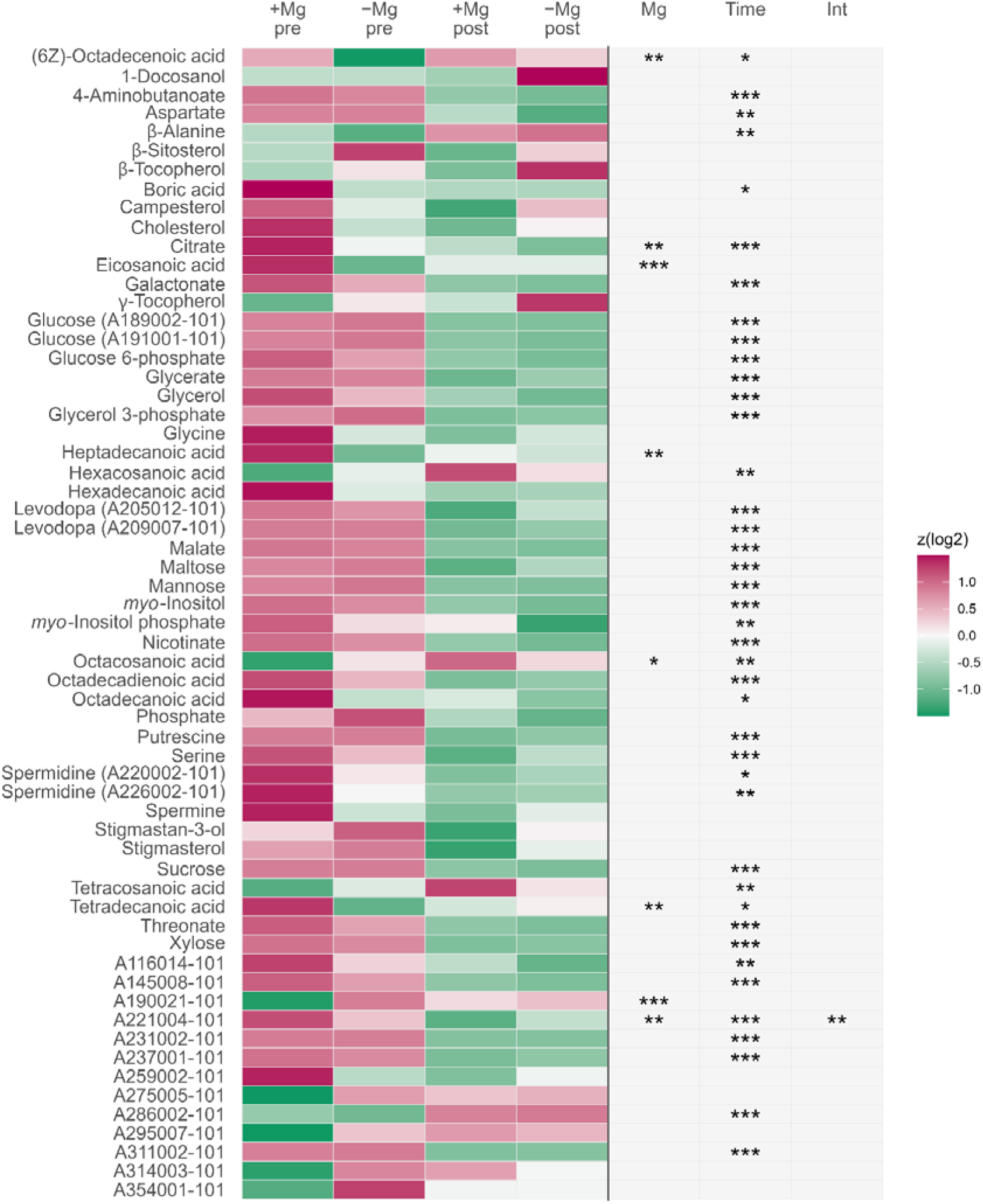
Heatmap of metabolome analysis of red light treated GC-enriched material. +Mg received a well supply of MgSO _4_ (0.3 m M MgSO4 ) in a hydroponic system, whereas no Mg was supplied in - Mg. GCs were isolated at two time points: pre-dawn and post-dawn after 2 h of red light (50 µmol m⁻² s⁻¹). Each metabolite is represented by four cells corresponding to the Mg × time-of-day combinations (+Mg and −Mg, pre- and post-dawn). Values were log2-transformed, and the mean of each group (n ≥ 4) was row-wise z-scored for display.F alse colors indicate the resulting z(log2) value according to the scale.. A two-way aligned rank transform (ART) ANOVA was applied and P-values were adjusted for multiple testing using the Benjamin-Hochberg false discovery rate (FDR) correction. The three columns on the right (Mg, Time, Int) give the adjusted significance of the Mg main effect (Mg), the time-of-day main effect (Time) and the Mg × time-of-day interaction (Int). Statistical significance is indicated as follows: *P < 0.05, **P < 0.01, ***P < 0.001. Compounds named by identifiers A######-101 indicate frequently observed but yet non-identified metabolites archived by the Golm Metabolome Databas e (http://gmd.mpimp-golm.mpg.de/). Created using RStudio.

As sampling time was the dominant source of variation in GC enriched material (Fig. 8), only the pre-dawn samples were statistically analyzed. The first component separated pre-from post-dawn samples, whereas the two Mg^2+^ treatments overlapped and the post-dawn groups converged irrespective of Mg^2+^ supply. In pre-dawn samples, the two Mg²⁺ treatments separated along the first component with a cross-validated BER of 0.15 (permutation P = 0.005).

**Figure 8:**
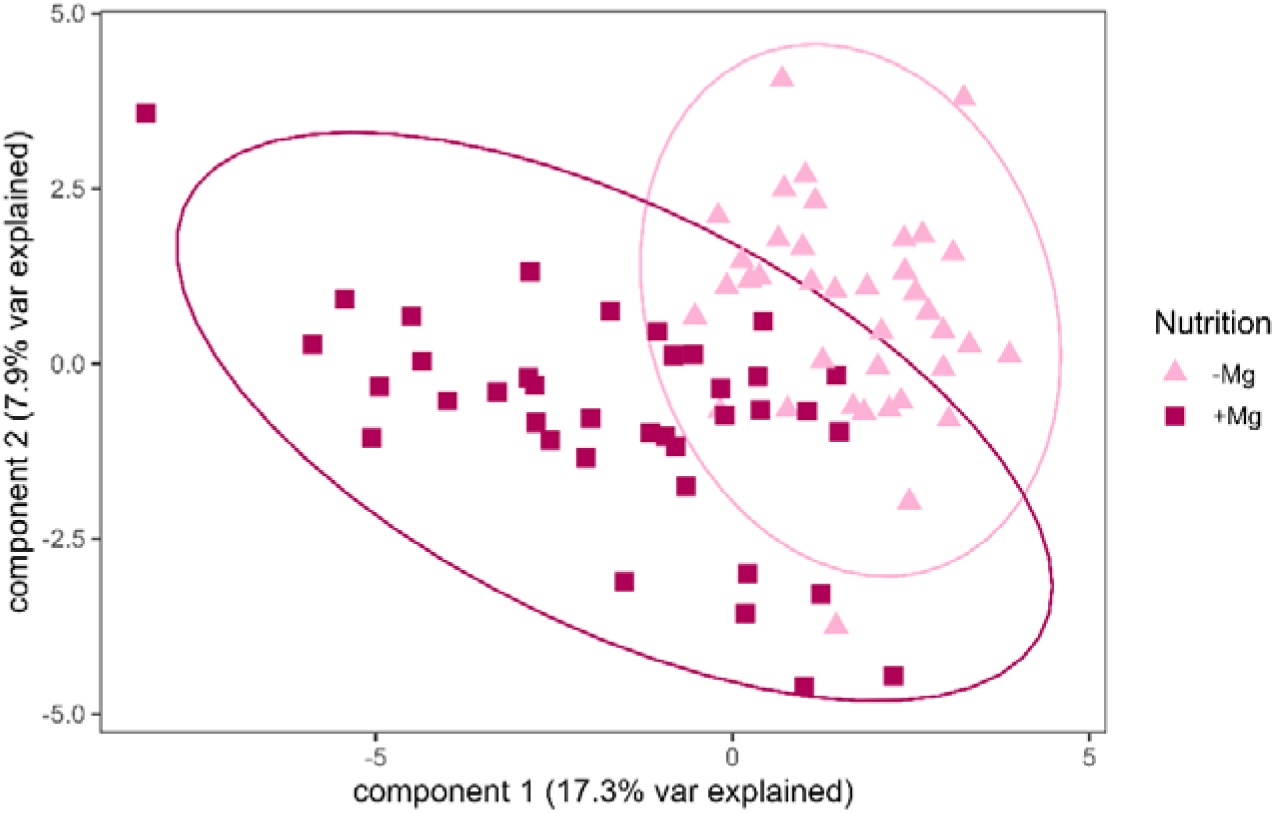
Partial Least Squares–Discriminant Analysis (PLS⍰DA) score plot of the GC metabolome showing sample separation. along component 1 (23.1% variance explained) and component 2 (15.0% variance explained). Points represent individual samples according to Mg²□ treatment (colour) and time of day (shape: circles, pre⍰dawn; triangles, post⍰dawn). Ellipses denote 95% confidence regions. Created using RStudio.

Regarding the proteomic profile in GCs, a total of 1708 proteins were identified of which 1091 had an annotated biological process. PLS-DA with 1 component explained 14.34% of variance in X and 100% in Y and separated the samples by Mg^2+^ treatment (Figure 9). Based on the VIP threshold, 277 proteins were included in GO enrichment analysis. Photosynthesis- and translation-related GO categories were enriched (p_adj_ < 0.05) (Table 1). All 38 significantly enriched proteins (Table S3) were more abundant in -Mg than +Mg treatment.

**Figure 9:**
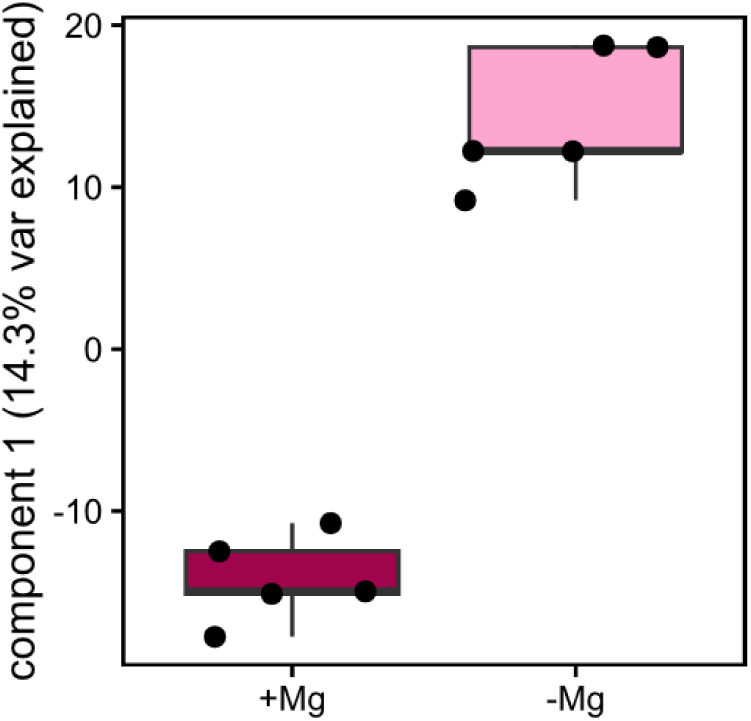
Partial Least Squares–Discriminant Analysis (PLS-DA) score plot showing proteome sample separation along component 1 (14.3% variance explained). Points represent individual samples according to Mg^2+^ treatment.

**Table 1:** Gene Ontology (GO) enrichment of Biological Process of proteins with VIP score higher than the breakpoint. Enrichment was performed using topGO (algorithm: weight01, Fisher’s exact test) with the BP-annotated protein universe (n = 1091). P-values were adjusted for multiple testing using Benjamini–Hochberg correction. Fold change represents the relative protein abundance of -Mg compared to +Mg.

| GO ID | GO Term | Background | Significant | Fold change | p<sub>adj</sub> |
| --- | --- | --- | --- | --- | --- |
| GO:0015979 | Photosynthesis | 46 | 13 | 3.80 | 0.02712 |
| GO:0006412 | Translation | 106 | 25 | 3.18 | 0.00086 |

## 4 Discussion

### 4.1 Pre-chlorotic Mg^2+^ deficiency reaches guard cells while leaves remain structurally intact

In vegetative shoot parts, plants require 1.5–3.5 g Mg^2+^ kg⁻¹ DM (equivalent to 0.15–0.35% DM) to secure optimal growth (P. Marschner, 2012). The exact critical Mg^2+^ threshold, however, varies between plant species, tissue, and developmental stage (Gransee & Führs, 2013). In order to have *V. faba* plants for this study that serve as a positive control with regard to Mg^2+^-nutrition, plants were cultivated with a well supply of 0.3 mM Mg^2+^ in the nutrient solution. These plants reached a leaf Mg^2+^ concentration of 0.31% DM in the second fully expanded leaves (Fig. 2A). Because the leaf lamina is composed predominantly of photosynthetically active mesophyll tissue (Lawson, 2009), bulk leaf Mg²⁺ concentration is considered a good proxy for the Mg²⁺ status of mesophyll cells, where some physiological processes analyzed in this study take place. In contrast, withholding Mg^2+^ from the nutrient solution significantly reduced the leaf Mg^2+^ concentration to 0.08% DM (Fig.2A). Thus, Mg^2+^ deficiency was established in these leaves. The severity of the Mg²⁺ deficiency was deliberately adjusted so that the leaves had not yet developed interveinal chlorosis (Fig. 1B). This ensured that the physiological measurements were performed in non-collapsed tissues and cells, allowing the primary effects of Mg²⁺ deficiency to be assessed. If the plants were grown for an additional 7 days, the typical Mg²⁺ deficiency symptoms would become visible, beginning with interveinal chlorosis followed by tissue necrosis (Fig. 1C).

Mg^2+^ homeostasis in guard cells contributes to stomatal opening (Inoue et al., 2022), but it remains unknown whether Mg^2+^ deficiency reaches GC before visible chlorosis. In our study, Mg^2+^ concentration in GC-enriched material decreased from 0.09% DM in Mg^2+^-sufficient plants to 0.04% DM under Mg^2+^ deficiency (Fig. 2B). To our knowledge, this is the first evidence that Mg^2+^ deficiency reduces Mg^2+^ concentration in GC-enriched material before visible chlorosis. Bulk-leaf Mg^2+^ declined more strongly than Mg^2+^ in the GC-enriched fraction. However, absolute concentrations between these fractions cannot be directly compared because their cellular composition differs. (Fig. 2). Notably, Mg^2+^ concentrations in GC-enriched material cannot be evaluated against established deficiency thresholds because such thresholds are based on bulk vegetative tissue. Notably, even the Mg^2+^-sufficient GCs fell below the optimal range defined for vegetative tissue. However, bulk-tissue Mg^2+^ thresholds cannot be directly applied to GC-enriched material. This is supported by Conn et al., 2011, who showed that Mg^2+^ is not evenly distributed among different cell types of Arabidopsis leaves, with mesophyll cells reaching the highest Mg^2+^ concentrations, while epidermal cells show lower concentrations. Thus, Mg^2+^ was clearly reduced in GC-enriched material, but this reduction cannot be related to a defined GC-specific deficiency threshold.

To verify that Mg^2+^ deficiency had not yet progressed to a stage of severe chlorotic or structurally compromised, i.e., heavily dysfunctional leaf tissue, we evaluated MCARI1 in the second fully expanded leaves. This index is mainly related to the green leaf area signal and is comparatively insensitive to pigment variation (Haboudane, 2004). MCARI1 did not differ significantly between Mg^2+^ treatments (Fig. 3A+D), neither on the abaxial nor on the adaxial leaf surface. This supports the view that the measured leaves still represented green, structurally intact tissue, although concentration of Mg^2+^ was lower than required. It is stated that an early consequence of Mg^2+^ deficiency is the accumulation of sucrose in leaves (Hermans et al., 2004). This carbohydrate accumulation decreases *A* due to negative feedback response. Together with the dependence of Rubisco activation on Mg²⁺, this results an imbalance between absorbed and utilised light energy. The generation of ROS is enhanced due to this resulting over-reduction of the photosynthetic electron transport chain, which photooxidatively degrade chlorophyll and thylakoid constituents (H. Marschner & Cakmak, 1989). Mg^2+^ functions as the central atom of chlorophyll and is therefore essential for photosynthesis (Z. C. Chen et al., 2018). As a consequence, Mg^2+^ deficiency may alter the pigment status, which can be detected through spectral reflectance. NDVI integrates information on both chlorophyll content and green leaf area. Since MCARI1 did not differ between treatments, the significant abaxial reduction in NDVI under Mg^2+^ (Fig. 3F) is unlikely the result of a change in leaf area and can instead be attributed primarily to a decline in chlorophyll on this surface. As the adaxial surface in dorsiventral dicotyledonous leaves, which includes faba bean, is primarily associated with light capture (Fukushima & Hasebe, 2014), these findings could reflect a preferential retention of Mg^2+^ and chlorophyll on this surface to sustain light harvesting. This is supported by Wall et al. (2022) who observed the adaxial side contributing more to photosynthesis than the abaxial side in some species. CRI1 indicates the carotenoid content (Gitelson et al., 2002) and was significantly higher on the adaxial side under Mg^2+^ deficiency, whereas no significant difference was detected on the abaxial side. This finding is discussed below in the context of photosynthetic limitation (see Section 4.2).

### 4.2 Guard cells are affected, but do not explain the loss of carbon gain

It was hypothesised that Mg^2+^ deficiency might affect E not only through impaired mesophyll photosynthesis, but also directly through Mg^2+^-dependent processes intrinsic to GCs. This distinction is important because a reduction in *E* can arise from partial stomatal closure, but stomatal closure may either be driven by mesophyll-derived signals or by functional changes intrinsic to the GCs. Under 1000 µmol m⁻² s⁻¹ red light, A was significantly reduced under Mg^2+^ deficiency in L2 leaves (Fig. 4C). *E* and g_s_ were also significantly reduced (Fig. 4D,E), indicating partial stomatal closure. Thus, Mg^2+^ deficiency affected both carbon gain and water loss, raising the question of whether the reduced *E* was primarily a consequence of mesophyll limitation, GC dysfunction, or both.

These results are consistent with findings in maize (Jezek et al., 2014), barley (Jamali Jaghdani et al., 2021) and citrus seedlings (Ye et al., 2019). A decreased *A* under Mg^2+^ deficiency could possibly be explained by a reduced Rubisco abundance and rubisco activity (Peng et al., 2015). Additionally, an accumulation of sucrose under Mg^2+^ deficiency was observed in leaves of bean plants (Cakmak et al., 1994) and sugar beet (Hermans et al., 2005), which can lead to a negative feedback on Rubisco (Tränkner & Jamali Jaghdani, 2019) and may be a possible driver of the impaired photosynthesis. As the proteome and metabolome data were obtained from GC-enriched material, their abundances at the mesophyll level cannot be addressed with our data. Notably, sucrose did not differ significantly between the Mg^2+^ treatments in GC-enriched material (Fig. 7). The sucrose accumulation reported in whole leaves under Mg^2+^ deficiency (Cakmak et al., 1994; Hermans et al., 2005) was therefore not reflected at the GC level. This may indicate that GCs respond differently from mesophyll cells. However, the whole-leaf accumulation arises from mesophyll photosynthesis and impaired phloem export, a context that is necessarily absent in isolated epidermal fragments incubated without mesophyll supply.

Also, the Ci did not decrease significantly in the Mg^2+^-deficient treatment but was maintained at control levels (Fig. 4F). A parallel decline in g_s_ and A could leave Ci unchanged (Fig. 4C, E, F), because a reduction in CO_2_ demand, which would raise Ci, could be offset by a decrease in stomatal conductance. A predominantly stomatal limitation would be expected to lower Ci (Flexas & Medrano, 2002). The unchanged Ci is therefore consistent with a substantial non-stomatal contribution to the reduction in A. This agrees with recent findings in rice, where Mg^2+^ deficiency reduced photosynthesis mainly through limitations in mesophyll conductance and biochemical capacity, together with a smaller contribution of g*s* (Zhou et al., 2024). However, a stomatal contribution cannot be excluded. Since Ci reflects the balance between CO_2_ supply and demand, a parallel decline in g_s_ and A can result in a Ci not differing from the control level. This is supported by findings of Flexas and Medrano, 2002 who demonstrate that stomatal and non-stomatal limitations under stress conditions can be present at the same time. Taken together, the impairment of photosynthesis under Mg^2+^ deficiency likely includes non-stomatal limitations.

A closer look to L2 leaves revealed that the Y(II) was significantly reduced under Mg^2+^ deficiency (Fig. 4A), indicating that absorbed light energy was less efficiently used for photochemical electron transport. Y(NPQ) did not differ significantly under Mg^2+^ deficiency (Fig. 4B). Thus, the reduced Y(II) was not accompanied by an enhanced dissipation of excess excitation energy as heat. CRI1, which is a reflectance index sensitive to carotenoid content, was significantly higher under Mg^2+^ deficiency in the adaxial side of the leaves (Fig. 3B). A higher carotenoid concentration under Mg^2+^ deficiency was also observed in citrus seedlings (Ye et al., 2019). As Y(NPQ) was not significantly enhanced, the indicated enlarged carotenoid pool in the leaves, points instead to an antioxidant role, which is consistent with the proposed lipid-protective antioxidant function demonstrated by Johnson et al., 2007. Carotenoids function as quenchers of triplet chlorophyll and singlet oxygen (Ramel et al., 2012). Such a role would be consistent with an increased ROS load, as the excess excitation energy arising from constrained photochemistry promotes the formation of ROS. Consistent with this, a meta-analysis conducted by Hauer-Jákli & Tränkner, 2019 revealed a significantly higher abundance of ROS, as well as higher concentrations of ROS scavenging enzymes under Mg^2+^ deficiency.

To move beyond whole-leaf responses and directly address whether Mg^2+^ deficiency also affects GC function in L2 leaves, we next used microscopy-PAM to quantify PSII performance at the level of individual GC pairs.

The significant reduction of Y(II) indicates that GC chloroplast photochemistry itself was also impaired under Mg^2+^ deficiency (Fig. 5A), rather than being only inferred from whole-leaf photosynthetic limitation. Thus, Mg^2+^ deficiency reached the GC level and affected GC photochemical function. GC chloroplasts have been proposed to contribute to stomatal regulation, although their functional role remains debated (Lawson, 2009). Coordinated changes in mesophyll and GC photosynthesis were also reported in antisense tobacco with reduced photosynthetic capacity (Lawson et al., 2008). However, GCs of Arabidopsis plants with a specific chlorophyll deficiency in this cell type retained their ABA- and CO_2_-induced closing responses, indicating that GC photosynthesis does not directly influence these responses (Azoulay-Shemer et al., 2015). Therefore, the impaired Y(II) of GCs does not prove that GC photochemical impairment caused stomatal closure. It rather shows that the reduction in g_s_ occurred in a situation where whole-leaf photosynthesis was limited and GC photosynthesis was affected at the same time, but the exact cause of the limited g_s_ remains unclear.

### 4.3 Guard cells show photochemical impairment under Mg -deficiency without broad shifts in the analysed metabolite abundances

Because microscopy-PAM revealed that Mg^2+^ deficiency affected PSII performance at the level of individual GC pairs, we next asked whether this GC-resolved photochemical effect was accompanied by detectable changes in the analysed GC metabolite pools. Given the Mg^2+^ dependence of central processes such as Rubisco activity (Z. C. Chen et al., 2018) and consequently carbon assimilation (Tränkner et al., 2018), we initially expected changes in metabolites linked to sugar and primary carbon metabolism. Instead, a Mg²⁺ main effect was resolved for only 8 of the 61 analysed metabolites, confined to a fatty-acid/citrate cluster (Fig. 7), which were reduced under Mg^2+^ deficiency. Consistent with this, a PLS-DA separated the two Mg²⁺ treatments in pre-dawn samples. Thus, Mg^2+^ deficiency altered a small, specific subset of GC metabolites, rather than producing a broad shift in primary metabolism.

In contrast to the largely unchanged analysed metabolite abundances, the protein data indicated a Mg^2+^-dependent response in the GC-enriched fraction, with higher abundance of proteins assigned to photosynthesis- and translation-related GO categories under pre-chlorotic Mg²⁺ deficiency and red-light conditions (Table 1). Mg^2+^-deficient GCs showed significantly reduced effective quantum yield of Photosystem II (Fig. 5A) and thus a lower photosynthetic efficiency. Proteins related to photosynthetic electron transport in photosystem II (Photosystem II CP43 reaction center protein and Photosystem II D2 protein), chlorophyll biosynthesis (NADPH-protochlorophyllide oxidoreductase) and light harvesting (Chlorophyll a-b binding protein, chloroplastic) were significantly enriched in Mg^2+^ deficient GC. This is compatible with an adjustment of the GC photosynthetic apparatus under Mg^2+^ deficiency, but it does not by itself demonstrate functional compensation. In addition to light capture and electron transfer, proteins related to the Calvin-cycle (Phosphoribulokinase, Ribulose bisphosphate carboxylase) and sugar metabolism (Fructose-1,6-bisphosphatase, Pyrophosphate-fructose 6-phosphate 1-phosphotransferase) were also enriched. At the whole-leaf level, Mg^2+^ deficiency was associated with reduced CO2 assimilation (Fig. 4C) and reduced effective quantum yield of Photosystem II (Fig. 4A), consistent with previous reports that Mg^2+^ deficiency can impair photosynthetic processes (Peng et al., 2015). The enrichment of photosynthesis- and sugar-metabolism-related proteins in the GC-enriched fraction therefore suggests that GCs respond to Mg^2+^ deficiency at the protein level, even though this response was not mirrored by broad significant changes in the analyzed GC metabolite abundances. Because metabolite profiling was performed on isolated GC-enriched epidermal material, metabolic interactions with the mesophyll were not captured and may partly explain this discrepancy. Whether these protein changes support GC function, compensate for reduced GC photochemical efficiency, or reflect another adjustment cannot be resolved from the present data.

Because light-induced stomatal opening depends on ATP-consuming processes, especially plasma membrane H^+^-ATPase activity and subsequent ion uptake, we also examined whether Mg^2+^ deficiency affected the GC ATP pool. The result was that GC ATP concentration did not differ significantly between Mg^2+^ treatments at either time point. However, at pre-dawn it was numerically lower under Mg^2+^ deficiency, although this difference was not significant. In both treatments, absolute ATP was markedly lower at post-dawn than at pre-dawn. Light-induced stomatal opening requires ATP for plasma membrane H+-ATPase activity and subsequent ion uptake (Shimazaki et al., 2007). However, without a time-matched dark control, the decrease in ATP cannot be assigned specifically to light-induced stomatal opening. Nevertheless, it has to be considered that the total abundance of Mg^2+^ in the tissue cannot be equated with the free, metabolically active Mg^2+^ pool, as large proportions are bound (Z. C. Chen et al., 2018). So, neither bulk-leaf nor GC Mg^2+^ concentration necessarily reflect the free cytosolic Mg^2+^ available to the H^+^-ATPase. Furthermore, the activity of the H^+^-ATPase is set primarily by post-translational regulation (Kinoshita and Shimazaki, 2002) and cannot be inferred from total ATP concentration alone. Our measurements thus document an altered Mg^2+^ status of the GC-enriched fraction but do not permit a direct conclusion about H^+^-ATPase activity or the functionally available Mg^2+^/MgATP pool. Taken together, the analysed GC metabolites and ATP data do not provide evidence for a broad metabolic impairment of GCs under Mg^2+^ deficiency, whereas microscopy-PAM and protein abundance data indicate that GCs were affected at the photochemical and protein level.

## 5. Conclusion

This study shows that pre-chlorotic Mg^2+^ deficiency reaches GC-enriched material and reduces Y(II) in individual GC pairs. Thus, GC photochemistry is already affected before visible chlorosis. This was accompanied by changes in a small group of metabolites and higher abundance of photosynthesis-and translation-related proteins in GC-enriched material. At the whole-leaf level, A, gs and E decreased, but the contributions of mesophyll-derived signals and intrinsic guard-cell responses cannot be separated by our data.

## Acknowledgement

We thank the Norddeutsche Pflanzenzucht (NPZ, Holtsee, Germany) for providing *Vicia faba* seeds. We are grateful to Prof. Dr. Heiko Mibus-Schoppe for providing access to the phytotron chambers, especially for the large-scale experiment, which enabled the accommodation of more than 250 hydroponic pots. We also thank Mr. Jochen Dreis and Mr. Heinz-Peter Pletz for their support and assistance with the phytotron facilities during the experiment. We thank Mrs. Kerstin Ziesen and Mr. Robert Kunz for their valuable technical support during the experiment. We further thank Mrs. Manuela Günther for her guidance and assistance with metabolite extraction, and Mr. Ralph Lehnart for measuring Mg^2+^ concentrations in leaves and GCs. We gratefully acknowledge Prof. Dr. Ralf Schweiggert for providing access to the ultrasonic homogenizer and enabling its use for this study. We Finally acknowledge funding by the German Research Foundation (DFG, Deutsche Forschungsgemeinschaft) under project number 471624304.

## Author Contribution

C-M.G. conceived and designed the study. A.W. performed the experiments, analyzed the data, and developed the figures. E.F. and S.C. measured and analyzed the proteome data and prepared the related text, tables, and figures. M.I. measured the pH data, E.v.R. and J.Ki. measured the ATP data. A.E. and J.Ko. measured the metabolites, provided standardized data, and wrote the related Methods section, whereas K.K. analyzed the metabolite data and generated the related figures. A.W., C-M.G., K.K. and B-L.F. performed the data interpretation. A.W. wrote the Introduction and Materials and Methods, K.K. wrote the Results, Discussion, and Conclusion, C-M.G. wrote the Abstract. A.W., C-M.G., E.F. and K.K. reviewed and edited the manuscript, with input from all authors. All authors reviewed and approved the final version.

## Supplementary material

### Supplemental figure

**Figure S1:**
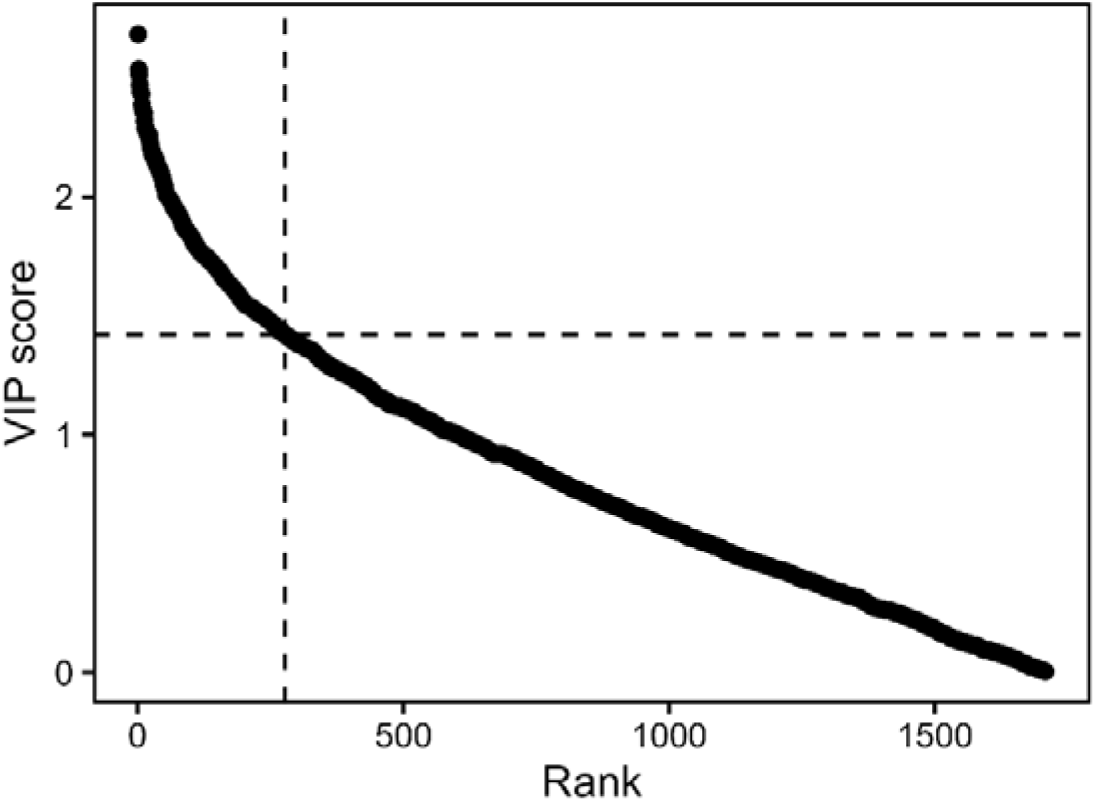
Variance Importance Projection (VIP) score versus rank of all 1708 identified proteins used in PLS-DA analysis. The dashed line represents the breakpoint at rank 278 with VIP score of 1.420543. Proteins with VIP higher than the breakpoint added explanatory value to separate -Mg and +Mg treated

### Supplemental tables

**Table S1:**
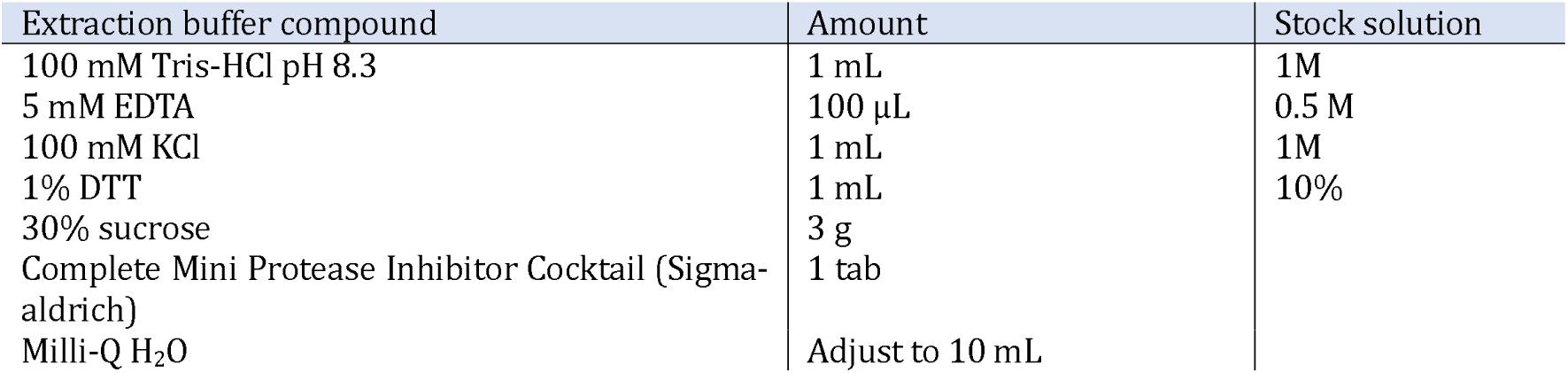
Extraction buffer composition. Amounts are given for a total volume of 10 mL.

**Table S2:** Lysis buffer composition. Amounts are given for a total volume of 20 mL.

| Lysis buffer compound | Amount |
| --- | --- |
| 8 M urea | 9.6 g |
| 5 mM DTT | 0.0154 g |
| 30 mM Tris | 0.0728 g |
| Milli-Q H<sub>2</sub>O | Adjust to 20 mL |

**Table S3:**
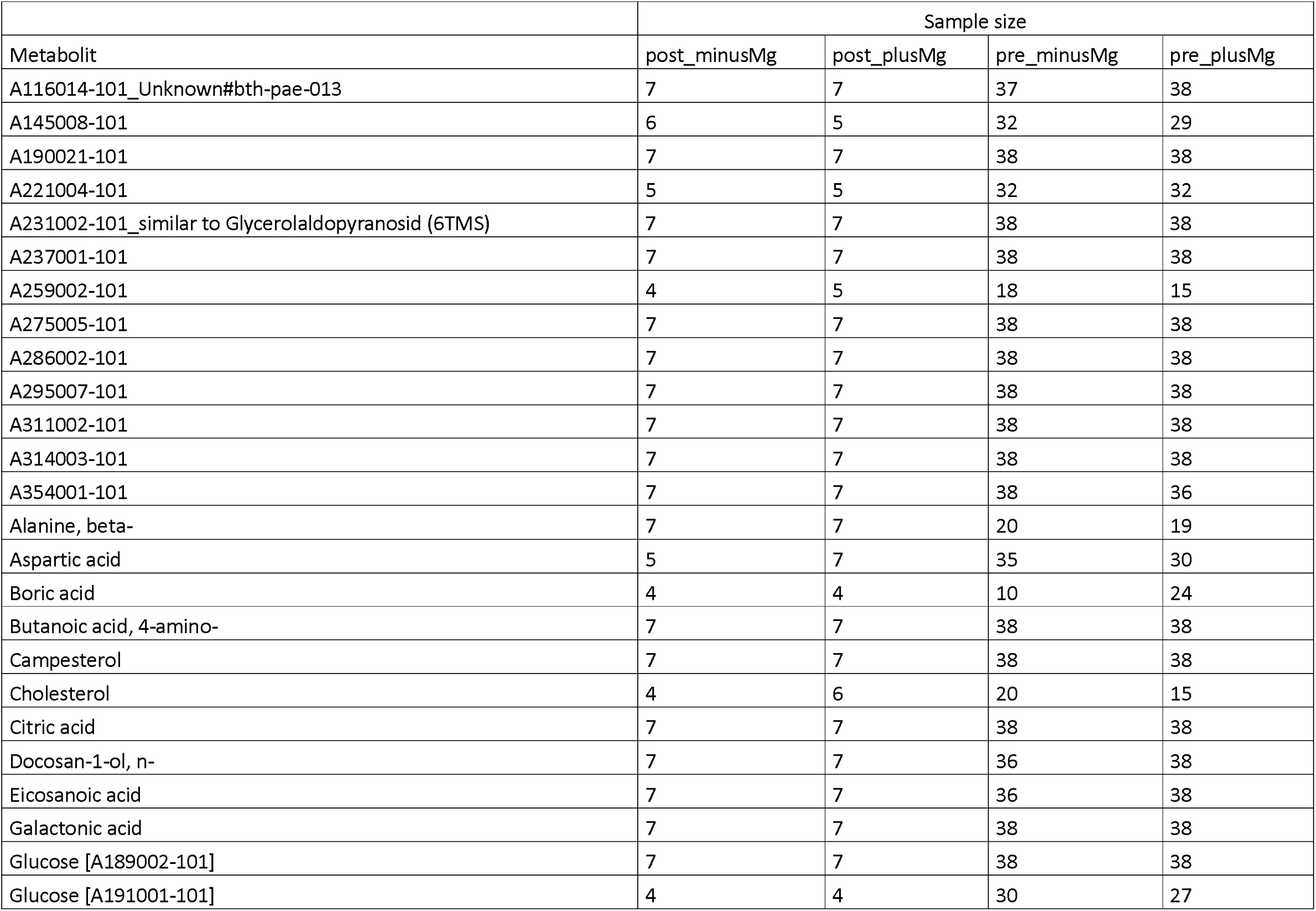
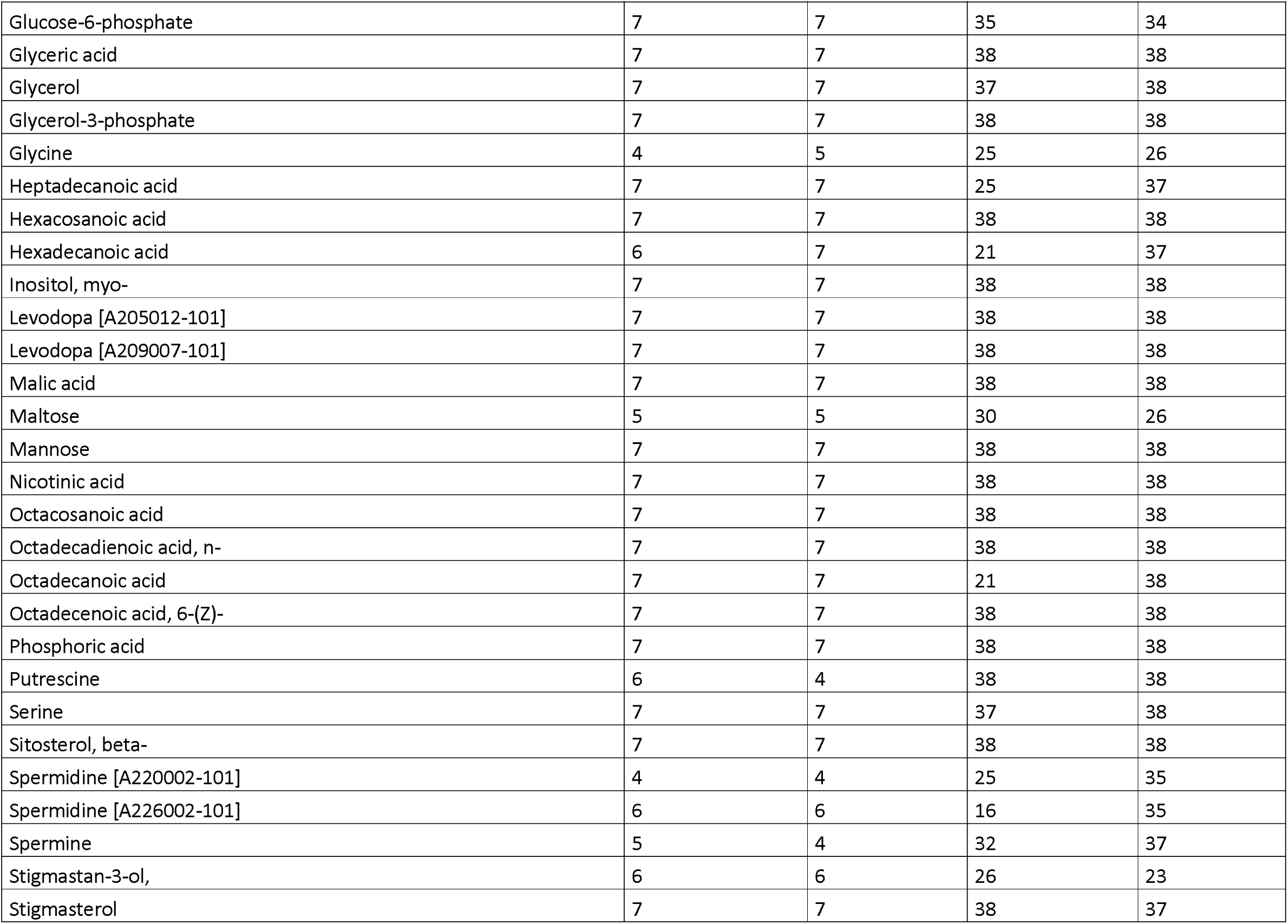
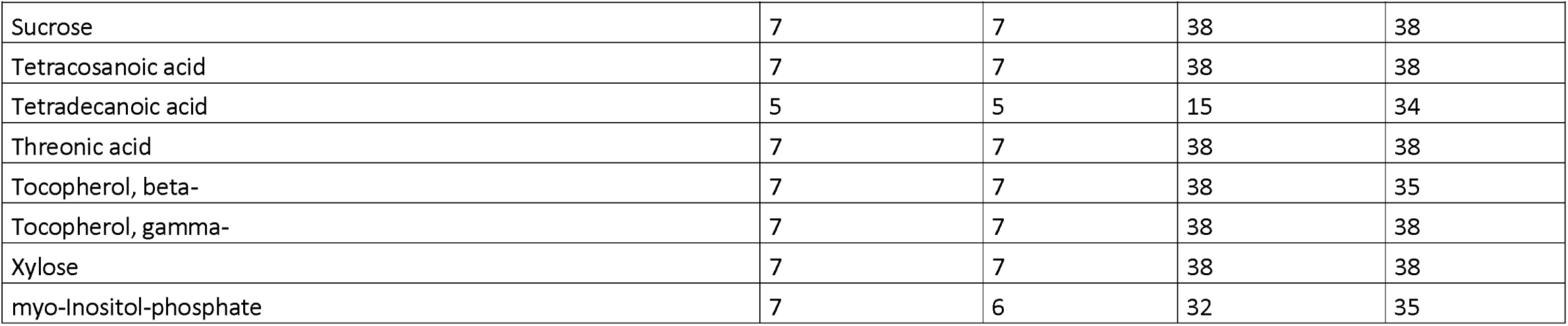
Sample size of the metabolites for each Mg^2+^ treatment and sample time. Plus_Mg = Mg^2+^-sufficient treatment, Minus_Mg = Mg^2+^-deficient treatment, post = post-dawn, pre = pre-dawn.

| Metabolit | Sample size |  |  |  |
| --- | --- | --- | --- | --- |
|  | post_minusMg | post_plusMg | pre_minusMg | pre_plusMg |
| A116014-101_Unknown#bth-pae-013 | 7 | 7 | 37 | 38 |
| A145008-101 | 6 | 5 | 32 | 29 |
| A190021-101 | 7 | 7 | 38 | 38 |
| A221004-101 | 5 | 5 | 32 | 32 |
| A231002-101_similar to Glycerolaldopyranosid (6TMS) | 7 | 7 | 38 | 38 |
| A237001-101 | 7 | 7 | 38 | 38 |
| A259002-101 | 4 | 5 | 18 | 15 |
| A275005-101 | 7 | 7 | 38 | 38 |
| A286002-101 | 7 | 7 | 38 | 38 |
| A295007-101 | 7 | 7 | 38 | 38 |
| A311002-101 | 7 | 7 | 38 | 38 |
| A314003-101 | 7 | 7 | 38 | 38 |
| A354001-101 | 7 | 7 | 38 | 36 |
| Alanine, beta- | 7 | 7 | 20 | 19 |
| Aspartic acid | 5 | 7 | 35 | 30 |
| Boric acid | 4 | 4 | 10 | 24 |
| Butanoic acid, 4-amino- | 7 | 7 | 38 | 38 |
| Campesterol | 7 | 7 | 38 | 38 |
| Cholesterol | 4 | 6 | 20 | 15 |
| Citric acid | 7 | 7 | 38 | 38 |
| Docosan-1-ol, n- | 7 | 7 | 36 | 38 |
| Eicosanoic acid | 7 | 7 | 36 | 38 |
| Galactonic acid | 7 | 7 | 38 | 38 |
| Glucose [A189002-101] | 7 | 7 | 38 | 38 |
| Glucose [A191001-101] | 4 | 4 | 30 | 27 |
| *Table S3 continued* |  |  |  |  |
| Glucose-6-phosphate | 7 | 7 | 35 | 34 |
| Glyceric acid | 7 | 7 | 38 | 38 |
| Glycerol | 7 | 7 | 37 | 38 |
| Glycerol-3-phosphate | 7 | 7 | 38 | 38 |
| Glycine | 4 | 5 | 25 | 26 |
| Heptadecanoic acid | 7 | 7 | 25 | 37 |
| Hexacosanoic acid | 7 | 7 | 38 | 38 |
| Hexadecanoic acid | 6 | 7 | 21 | 37 |
| Inositol, myo- | 7 | 7 | 38 | 38 |
| Levodopa [A205012-101] | 7 | 7 | 38 | 38 |
| Levodopa [A209007-101] | 7 | 7 | 38 | 38 |
| Malic acid | 7 | 7 | 38 | 38 |
| Maltose | 5 | 5 | 30 | 26 |
| Mannose | 7 | 7 | 38 | 38 |
| Nicotinic acid | 7 | 7 | 38 | 38 |
| Octacosanoic acid | 7 | 7 | 38 | 38 |
| Octadecadienoic acid, n- | 7 | 7 | 38 | 38 |
| Octadecanoic acid | 7 | 7 | 21 | 38 |
| Octadecenoic acid, 6-(Z)- | 7 | 7 | 38 | 38 |
| Phosphoric acid | 7 | 7 | 38 | 38 |
| Putrescine | 6 | 4 | 38 | 38 |
| Serine | 7 | 7 | 37 | 38 |
| Sitosterol, beta- | 7 | 7 | 38 | 38 |
| Spermidine [A220002-101] | 4 | 4 | 25 | 35 |
| Spermidine [A226002-101] | 6 | 6 | 16 | 35 |
| Spermine | 5 | 4 | 32 | 37 |
| Stigmastan-3-ol, | 6 | 6 | 26 | 23 |
| Stigmasterol | 7 | 7 | 38 | 37 |
| *Table S3 continued* |  |  |  |  |
| Sucrose | 7 | 7 | 38 | 38 |
| Tetracosanoic acid | 7 | 7 | 38 | 38 |
| Tetradecanoic acid | 5 | 5 | 15 | 34 |
| Threonic acid | 7 | 7 | 38 | 38 |
| Tocopherol, beta- | 7 | 7 | 38 | 35 |
| Tocopherol, gamma- | 7 | 7 | 38 | 38 |
| Xylose | 7 | 7 | 38 | 38 |
| myo-Inositol-phosphate | 7 | 6 | 32 | 35 |

**Table S4:**
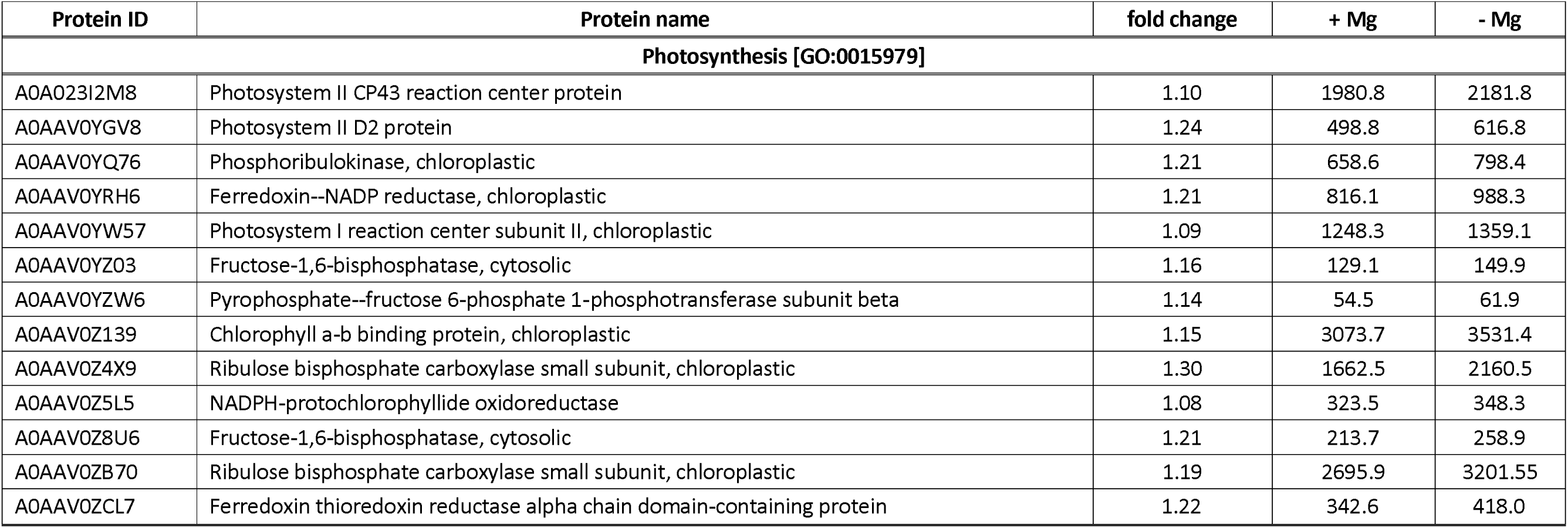
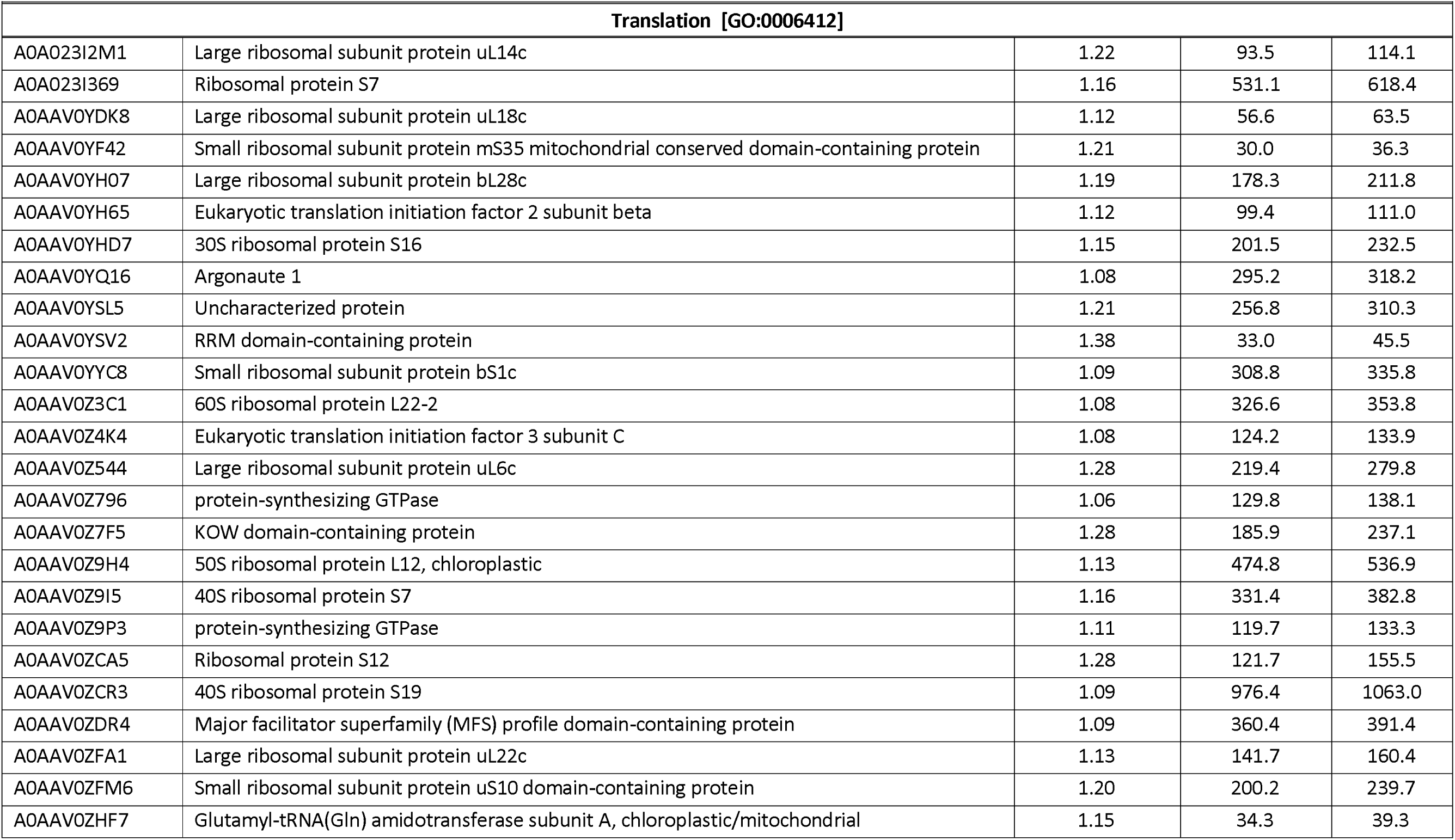
Significantly enriched proteins according to Gene Ontology analysis related to photosynthesis and translation processes. Fold change represents the relative protein abundance of - Mg compared to +Mg.

**Supplemental Table S5:** Raw and log2-transformed metabolite data from GC-enriched material are provided as a separate downloadable file.

## Data Availability Statement

Data will be made available on request.

